# Non-microglial downregulation of *PLCG2* impairs synaptic function and elicits Alzheimer disease-related hallmarks

**DOI:** 10.1101/2024.04.29.591575

**Authors:** Audrey Coulon, Florian Rabiller, Mari Takalo, Avishek Roy, Alexandre Pelletier, Henna Martiskainen, Dolores Siedlecki-Wullich, Nina Lannette-Weimann, Naďa Majerníková, Arthur Grenon, Vance Gao, Anaël Erhardt, Anne Pernodet, Morgane Lemaire, Floriane Limoge, Pauline Walle, Tiago Mendes, Karine Guyot, Célia Lemeu, Lukas-Iohan Carvalho, Ana Raquel Melo de Farias, Marc Hulsman, Chloé Najdek, Alejandra Freire-Regatillo, Orthis Saha, Philippe Amouyel, Camille Charbonnier, Jean-François Deleuze, Orio Dols-Icardo, Heli Jeskanen, Roosa-Maria Willman, Teemu Kuulasmaa, Mitja Kurki, John Hardy, Sami Heikkinen, Henne Holstege, Petra Mäkinen, Gaël Nicolas, Simon Mead, Michael Wagner, Alfredo Ramirez, Tuomas Rauramaa, Aarno Palotie, Rebecca Sims, Hilkka Soininen, John van Swieten, Julie Williams, Céline Bellenguez, Carla Gelle, Erwan Lambert, Marcos R. Costa, Julia TCW, Enrico Glaab, Anne-Marie Ayral, Florie Demiautte, Benjamin Grenier-Boley, Manon Muntaner, Delphine Eberlé, Séverine Deforges, Joel Haas, Devrim Kilinc, Christophe Mulle, Julien Chapuis, Mikko Hiltunen, Julie Dumont, Jean-Charles Lambert

## Abstract

We developed a high content screening to investigate how Alzheimer disease (AD) genetic risk factors may affect synaptic mechanisms in rat primary neuronal cultures. Out of the target genes identified, we found that *Plcg2* downregulation in mouse dentate gyrus neurons consistently disrupted dendritic morphology and synaptic function. In human neuronal cultures (hNCs), *PLCG2* downregulation also impaired synaptic function and increased Aβ levels and Tau phosphorylation. Very rare *PLCG2* loss-of-function (LoF) variants were associated with a 10-fold increased AD risk. *PLCG2* LoF carriers exhibit low mRNA/protein *PLCG2/*PLCγ2 levels and the R953* LoF mutation compromised synaptic function and increased AD hallmarks in hNCs. Single nuclei RNAseq analyses confirmed that the downregulation of *PLCG2* impacted pathways related to synaptic and neuronal functions, potentially through neurexin in neurons. In conclusion, PLCγ2 downregulation could increase AD risk by impairing synaptic functions and increasing the Aβ levels and Tau phosphorylation in neurons.

## INTRODUCTION

Alzheimer’s disease (AD) is characterized by intracellular aggregation of abnormally hyperphosphorylated Tau protein and extracellular accumulation of amyloid-beta (Aβ) in plaques. According to the “amyloid cascade hypothesis”, Aβ aggregation is an early toxic condition that can induce Tau aggregation and neuronal death. However, while recent clinical studies using anti-Aβ immunotherapies in principle support this hypothesis^1^, it has been regularly revised to reflect the advancements in our understanding of AD^2–4^. It is now well accepted that the amyloid cascade hypothesis appears to be over-simplistic and fails to account for the intricacy and heterogeneity of the pathophysiological processes involved in the common forms of the disease.

Since the proportion of risk attributable to genetic susceptibility factors for AD has been estimated to be between 60% and 80% in twin studies^5^, defining the AD genetic component should help in better understanding the pathophysiological processes and in generating complementary or alternative hypotheses. The development of genome-wide association studies (GWAS) and high-throughput sequencing has considerably advanced the AD genetic landscape over the last fifteen years, leading to the characterization of 76 loci associated with the AD risk^6–8^. Based on this new genetic landscape of AD, and beyond inflammation processes^9^, we proposed synaptic fragility/sensitivity as a pathological trigger^10^, depending on genetic risk factors, some of which have already been linked to synaptic mechanisms, such as *BIN1*, *FERMT2*, *PTK2B* or *CD2AP*^9–12^. This notion fits with key clinical observations suggesting that synapse dysfunction/loss is (i) one of the earliest pathological hallmarks in the brain and (ii) the strongest marker of cognitive decline^13–15^. Within this background, we systematically addressed the potential implication of the genes located in the loci associated with AD risk (as reported in^6^) in synaptic mechanisms. For this purpose, we developed a high-content screening (HCS) approach to systematically analyze the impact of the downregulation of a large number of AD risk genes on synaptic density and to further explore the potential implication of these genes in synaptic function.

## RESULTS

### HCS identifies GWAS-defined genes as potential modulators of synapse density

We developed an HCS approach to systematically characterize the effect on synaptic density of 198 genes located in the 76 loci associated with AD risk and selected as being expressed in brain cells (according to publicly available RNAseq datasets, see Methods online). A lentiviral shRNA library targeting these 198 genes was developed. The screen was performed in 384-well plates using rat hippocampal primary neuronal cultures (PNCs) transduced at day 1 in vitro (DIV1) using two multiplicities of infection (MOI 2 and 4, **Fig. 1a,b**). Immunocytochemistry was conducted at DIV21 against Synaptophysin 1 (Syp), Homer1 (Hom1) and Map2, pre-synaptic, post-synaptic and somatodendritic markers, respectively (**Fig. 1c,d**; supplementary. Fig. S1a)^16^. Synaptic density was then assessed *via* a high-content analysis workflow by assigning each post-synaptic structure to the nearest pre-synaptic structure, as previously described^17^. Non-targeting shRNA (shNT) and shRNA against Syp (shSyp) were used as controls. The complete HCS results are presented in supplementary Table S1. We selected AD risk genes with the strongest effect on synaptic density (top and bottom 2.5%) in three independent experiments, resulting in five genes (*Usp6nl*, *Psmc3*, *Plcg2*, *Csnk1g1*, *Cyb561*) associated with lower synaptic density and four genes (*Snx1*, *Ica1l*, *Oplah* and *Nck2*) associated with higher synaptic density when downregulated (**Fig. 1c**). For these nine genes, we confirmed the impact of shRNA-mediated downregulation on protein levels in PNCs (supplementary Fig. S1b, c). In conclusion, our HCS approach allowed us to identify several genes located within the loci associated with AD risk that strongly modulate synaptic density in rat PNCs.

**Figure 1.**
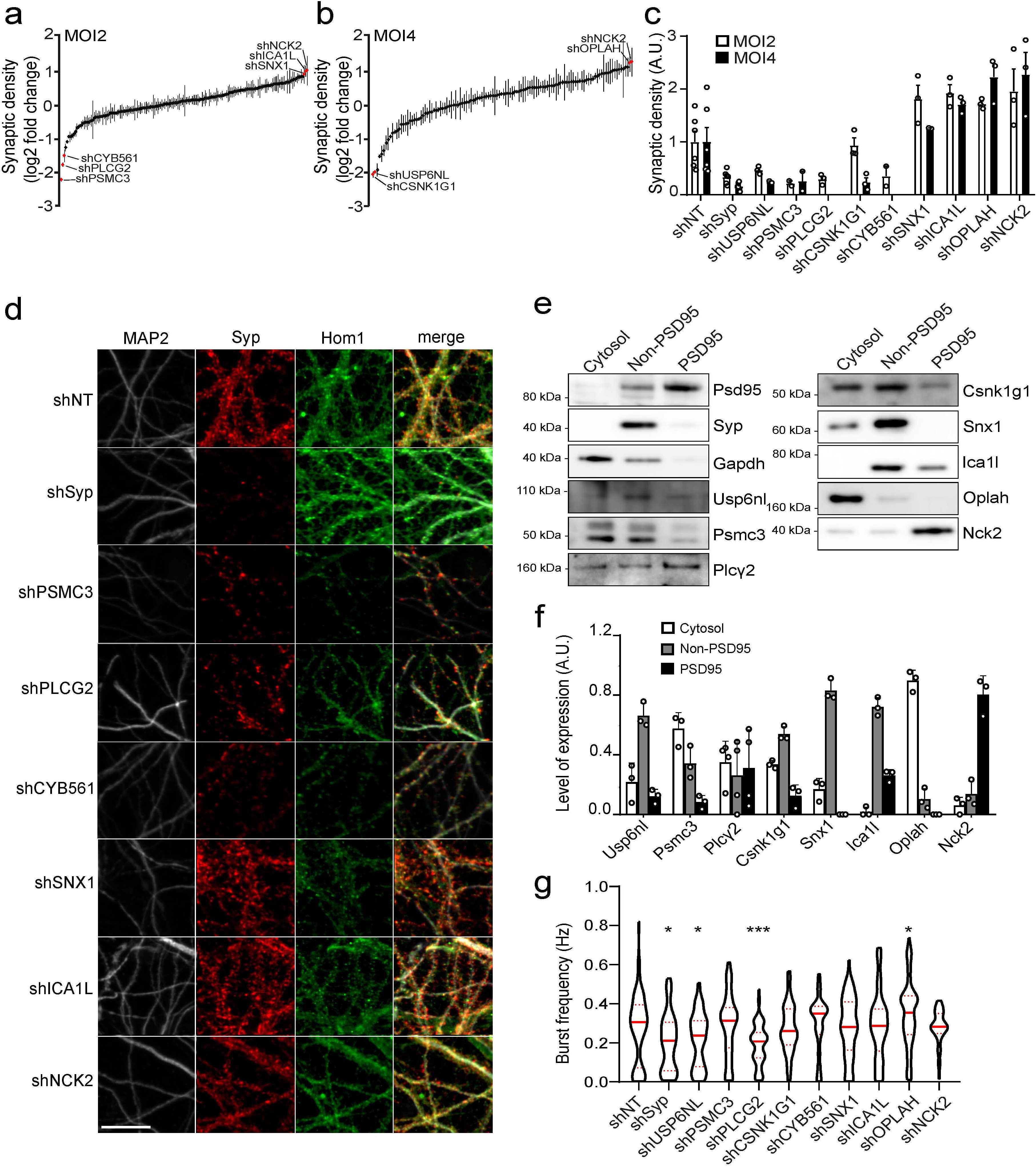
High-content screening identified nine AD-associated genes that significantly impact synaptic density and function in rat primary neuronal cultures. (a-b) Impact of shRNA-mediated downregulation of 198 AD-associated genes on synaptic density in rat hippocampal primary neuronal cultures (PNCs), virally transduced at MOI 2 (a) and MOI 4 (b). Data given as mean±SEM, n=3, top and bottom 2.5% genes are shown in red. (c) Quantification of synapse density following the assignment of Homer1 to Syp spots on Map2-positive dendrites. Data given as mean±SEM, n=3. (d) Representative images of Map2, Syp and Homer1 immunostainings following shRNA-mediated downregulation of *Syp, Psmc3, Plcg2, Cyb561, Snx1, Ica1l,* and *Nck2* at MOI 2; scale bar = 20 µm. (e-f) Immunoblotting (e) and quantification (f) of PSD and non-PSD fractions in primary rat hippocampal neuron synaptosomes (at DIV21) showing that Usp6nl, Psmc3, Plcγ2, Csnk1g1, Snx1, Ica1l, Oplah, and Nck2 proteins are expressed in the PSD95 and/or non-PSD95 fractions. Data given as mean±SD. (g) Impact of shRNA-mediated downregulation of *Syp, Usp6nl, Psmc3, Plcg2, Cyb561, Csnk1g1, Snx1, Ica1l, Oplah,* and *Nck2* on the burst frequency in PNCs at DIV21, measured by MEA. Violin plots show median and quartile lines, n=3. Statistical analyses were performed using Kruskal-Wallis test followed by Dunn’s multiple comparisons test. * p<0.05, *** p<0.001.

### *Plcg2* impacts electrophysiological properties of PNCs

We first determined whether proteins encoded by the nine selected genes could be detected in synaptosomes purified from PNCs (**Fig. 1e, f**). Except for Cyb561 (data not shown), we detected their presence in the PSD95 and/or non-PSD95 fractions. We then analyzed the impact of their downregulation on the electrophysiological properties of mature PNCs (at DIV21) using multielectrode arrays (MEAs). As a positive control, the downregulation of *Syp* significantly decreased burst frequency, a read-out for neuronal activity (**Fig. 1g**). Similarly, the downregulation of *Usp6nl* and *Plcg2* significantly decreased burst frequency whereas the opposite was observed upon *Oplah* downregulation. Of note, *Usp6nl* downregulation also significantly impacted mean spike frequency (supplementary. Fig. S1d). We further focused on *Plcg2* as this AD risk gene was believed to be almost exclusively expressed in microglia and, as a consequence, it has been extensively investigated in this cell type^18–21^. However, *PLCG2* is also expressed in glutamatergic neurons as evidenced in publicly available single nuclei RNA sequencing (sn-RNA seq) datasets from human brain^22,23^ (supplementary Fig. S2) and therefore merits further investigation of its possible role in synaptic function in the context of AD pathogenesis.

### *Plcg2* downregulation in the mouse dentate gyrus reduces dendritic arborization complexity and spine density of granule cells

*Plcg2/PLCG2* is expressed in the mouse and human dentate gyrus (DG)^24^, which is the main gateway from the entorhinal cortex to the hippocampus. Synaptic plasticity in the DG is strongly affected in mouse models of AD^25,26^. Lentiviral vectors encoding either a shRNA against *Plcg2* (LV-shPLCG2) or a non-targeting control shRNA (LV-shNT), combined with GFP were injected into the dorsal hippocampus of adult mice of both sexes in the ipsi-(LV-shPLCG2) and contra-lateral (LV-shNT) hemispheres, respectively (**Fig. 2a** and supplementary Fig. S3a, b). We stained brain sections with an antibody against IBA1, a marker for microglia cells (supplementary Fig. S3c) and found that the number of GFP expressing cells positive for IBA1 was negligible (for both LV-shPLCG2 and LV-shNT vectors), indicating that microglial cells were not transduced by the lentiviral vectors. Analysis of GFP fluorescence images of DG neurons from shNT (n=35 neurons) and shPLCG2 (n=42 neurons) injected mice (**Fig. 2b**) revealed that *Plcg2* downregulation resulted in a reduction in dendritic length (shNT = 148±12 µm *vs* shPLCG2 = 77±9 µm, p<0.001; **Fig. 2c**) and dendritic volume (shNT = 270±32 µm^3^ *vs* shPLCG2 = 87±5 µm^3^, p<0.001; **Fig. 2d**), but not in the complexity index (p=0.24) or total branch number (p=0.52; **Fig. 2e, f**). We observed a decrease in the number of intersections in a Sholl analysis (p<0.0001, intersection *vs* distance from soma F_(24, 1560)_=23.23; **Fig. 2g, h**), in the spine density (number of spines per µm: shNT = 0.70±0.04, n=73 neurites *vs* shPLCG2 = 0.27±0.01, n=75 neurites, p<0.0001; **Fig. 2i**) and alterations in the morphology of dendritic spines in shPLCG2-transduced DG granule cells, with decreased spine length (shNT = 0.73±0.06 µm, n=89 *vs* shPLCG2 = 0.53±0.05 µm, n=101, p<0.0001) and head diameter (shNT = 0.48±0.02 µm, n=100 *vs* shPLCG2 = 0.29±0.01 µm, n=103, p<0.0001; **Fig. 2j, k**). Overall, in congruence with our findings *in vitro*, *Plcg2* downregulation markedly impairs dendritic morphology and spine density in DG granule cells *in situ*.

**Figure 2.**
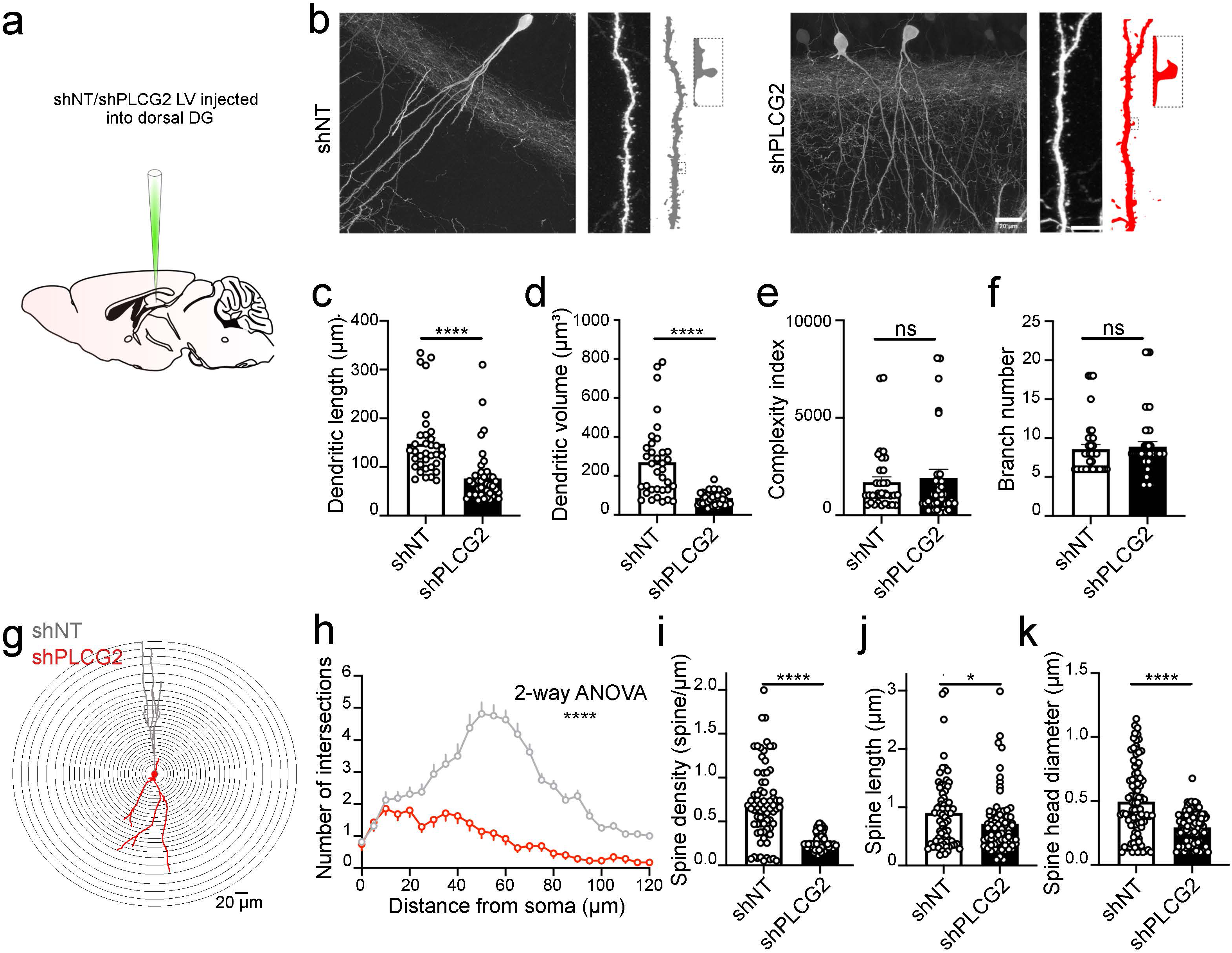
*Plcg2* downregulation alters the morphology of DG granule cells. (a) Schematic representation of the experimental approach to downregulate *Plcg2* in the DG. The lentiviral constructs were stereotaxically injected in either the ipsi-(LV-shPLCG2 for knock-down of *Plcg2*) or contralateral (LV-shNT, control) hemispheres. (b) High magnification immunofluorescence images of transduced DG cells as indicated by GFP expression, as well as a portion of dendritic branch with spines; scale bars = 20 µm for left panels and 7 µm for right panels. (c-f) Measurements of dendritic length (c), dendritic volume (d), complexity index (e), and branch number (f) in transduced DG granule cells. (g) Schematic diagram of the Sholl analysis, illustrating the morphological change in the dendritic tree of representative control (grey) *vs* shPLCG2 (red) transduced DG granule cells. Scale bar = 20 µm. (h) Distribution of intersections based on the Sholl analysis. (i-k) Measurements of average spine density (i), spine length (j) and spine head diameter (k), indicating a marked alteration in synaptic contacts onto DG granule cells. Data expressed as mean±SEM for bar graphs with individual data points. Sholl statistical analysis (n ≥90 per condition) was performed using two-way ANOVA test. All other statistical analyses were performed using Mann-Whitney test. * p<0.05, *** p<0.001, **** p<0.0001.

### *Plcg2* downregulation in the mouse dentate gyrus impairs electrophysiological properties of granule cells

We then prepared slices for the electrophysiological recording of DG granule cells transduced with shPLCG2 or shNT as visualized by their GFP fluorescence (**Fig. 3a**). We assessed the intrinsic electrophysiological properties (shNT, n=15 cells; shPLCG2, n=13 cells; **Fig. 3b, c** and supplementary Fig. S4a,b). *Plcg2* downregulation did not change the resting membrane potential (RMP) (shNT = – 57.2±2.2 mV *vs* shPLCG2 = –62.4±2.2 mV, p=0.11; **Fig. 3d**), but led to a reduction in the input resistance (shNT = 570±79 MOhm *vs* shPLCG2 = 312±35 MOhm, p=0.0061; **Fig. 3e**), which was accompanied by an increase in the half-width of action potentials (shNT = 1.17 ms *vs* shPLCG2 = 1.25 ms, p=0.027, supplementary Fig. S4c,d). We found that the amount of current required for the cell to fire an action potential (the rheobase) was significantly higher following *Plcg2* downregulation (shNT = –57±11 pA *vs* shPLCG2 = –86±10 pA, p=0.022; **Fig. 3f**). The number of spikes triggered by depolarizing pulses as a function of the amplitude of current injected also showed that shPLCG2-transduced DG neurons were less excitable (F_(1, 588)_ = 10.87, df=1, p=0.001, ordinary 2-way ANOVA; **Fig. 3g**). Overall, *Plcg2* downregulation leads to a substantial decrease in input resistance and in neuronal excitability. It can be suggested that Plcg2 contributes to the regulation of voltage-dependent ion channels around resting membrane potential of DG granule cells. We then performed voltage-clamp recordings in the presence of tetrodotoxin (TTX; 500 nM) to assess the impact of *Plcg2* downregulation on spontaneous miniature excitatory post-synaptic currents (mEPSCs) (**Fig. 3h**). We found significant differences in the average frequency of mEPSCs between shNT (n=27) and shPLCG2 (n=15) transduced neurons (shNT = 0.47±0.07 Hz *vs* shPLCG2 = 0.27±0.08 Hz, p=0.026; **Fig. 3i**). This result is consistent with the observed reduction of spine density, albeit to a lesser extent (–43% for mEPSC frequency *vs* –61% for spine density), which might be due to a compensatory increase in spontaneous release at the remaining synapses. In addition, *Plcg2* downregulation resulted in a decrease in the amplitude of mEPSCs (shNT = 13.6±0.7 pA *vs* shPLCG2 = 10.8±0.5 pA, p=0.034; cumulative distribution of mEPSC amplitudes, p=0.0008; **Fig. 3j, k**). We have finally recorded EPSCs evoked by the electrical stimulation of the perforant path in the presence of bicuculline to block GABA-A receptors (**Fig. 3l**). We measured the amplitude of AMPA-EPSCs (at –70 mV; **Fig. 3m**) and NMDA-EPSCs (at +40 mV in the presence of NBQX; **Fig. 3n**) and found that the NMDA/AMPA-EPSC ratio was largely reduced following *Plcg2* downregulation (shNT = 1.15, n=12 *vs* shPLCG2 = 0.31, n=13, p=0.019; **Fig. 3o**). Our data suggest that the downregulation of Plcg2 selectively decreases the amount of post-synaptic NMDARs at synapses in DG cells. In conclusion, in accordance with our results *in vitro*, *Plcg2* downregulation markedly impairs electrophysiological properties of DG granule cells.

**Figure 3.**
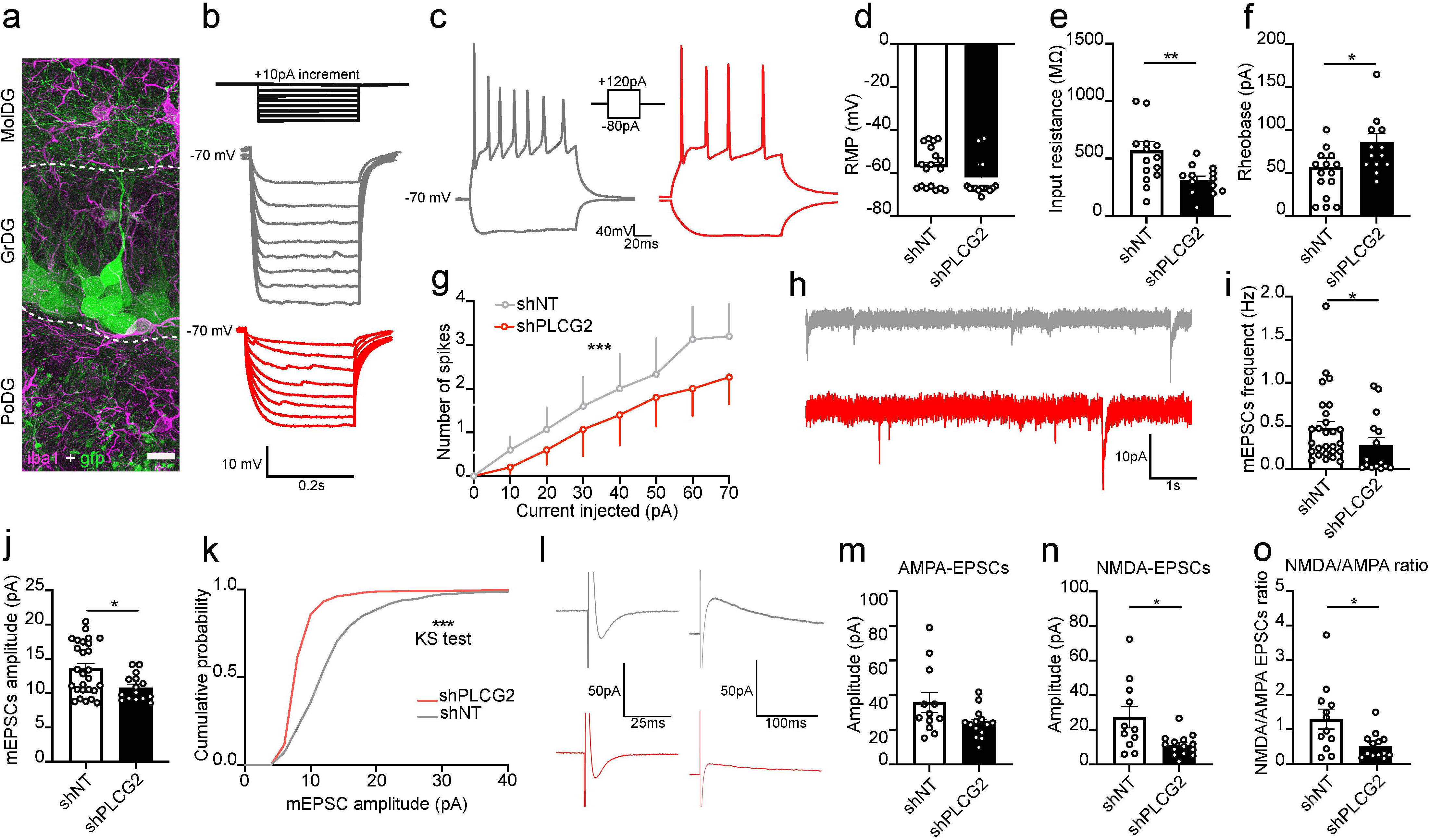
*Plcg2* downregulation in DG granule cells decreases neuronal excitability and impacts synaptic function. (a) Representative image of maximum intensity projection of section labeled with IBA1 (in magenta) and GFP (in green) showing the selective expression of viral vector in DG granule cells. MolDG= molecular layer of DG, GrDG= granular DG, PoDG= polymorph layer of DG. Patch-clamp recordings were performed from GFP expressing granule cells identified under the electrophysiology microscope. Scale bar = 10 µm. (b) Representative traces from shNT (in grey) and shPLCG2 (in red) transduced DG granule cells recorded in the current-clamp mode, with injection of current steps from –80 pA to 0 pA with 10 pA increments. (c) Representative traces of current-clamp recordings showing spike discharges triggered by a depolarization pulse of current (120 pA) from shNT (in grey) and shPLCG2 (in red) transduced DG cells. (d-f) Current-clamp recordings were used to assess resting membrane potential (RMP) (d), input resistance (e) and rheobase (f). (g) Number of spikes plotted against current injected. (h) Representative traces of miniature excitatory post-synaptic currents (mEPSCs) recorded in the voltage-clamp mode from shNT (in grey) and shPLCG2 (in red) DG cells in presence of 500 nM TTX. (i-k) Measurements of average frequency of mEPSCs (i), average amplitude of mEPSCs (j) and cumulative distribution of mEPSC amplitudes (k). (l) Representative traces of AMPA-EPSCs (at –70mV) and NMDA-EPSCs (at +40mV in the presence of 10 µM NBQX) evoked by the electrical stimulation of the perforant path (at 0.1 Hz). (m-o). Graphs representing the peak amplitude of AMPA-EPSCs (m) and of NMDA-EPSCs (n), and the ratio of NMDA-EPSCs *vs* AMPA-EPSCs (o) for each recorded neuron. Data are represented as mean±SEM with bar and individual data points for average values. All statistical analyses were performed using Mann-Whitney test with the exception of the cumulative distribution where the Kolmogorov-Smirnov (KS) test was applied. * p<0.05, ** p<0.01, ***p<0.001.

### Downregulation of *PLCG2* in human neuronal cultures negatively impacts synaptic density and function

We then sought to extend our findings in rodents to an *in vitro* model of human neurons, derived from human induced pluripotent stem cells (iPSCs, ASE-9109 cell line^27^). After 4 weeks of spontaneous differentiation from neural progenitor cells transduced with either shNT or shPLCG2, we obtained a mixed culture of mature human induced neurons and astrocytes, which we will refer to as human neuronal cultures (shNT-hNCs or shPLCG2-hNCs; **Fig. 4a**). We validated *via* immunoblotting that PLCγ2 downregulation in this mixed culture was specific as it did not impact PLCγ1 expression (**Fig. 4b**). To characterize the impact of *PLCG2* downregulation on synaptic density in hNCs, we used custom microfluidic devices in which mature neuronal synapses can be fluidically isolated in a distinct (synaptic) chamber and easily quantified. As in the case of HCS, we performed immunocytochemistry against synaptic markers and assessed synaptic density using a previously described algorithm^17^ (**Fig. 4c**). We fully replicated in hNCs the results obtained in the HCS experiments using rat NPCs, *i.e.*, *PLCG2* downregulation decreased synaptic density (**Fig. 4c,d**). Similar results were obtained when using rat PNCs cultured in microfluidic devices (supplementary Fig. S5). Lastly, we analyzed the impact of *PLCG2* downregulation on the electrophysiological properties of hNCs using MEAs. Again, we replicated in hNCs the results we obtained in rat PNCs, *i.e.*, *PLCG2* downregulation induced a significant decrease in burst frequency, and a downward trend was observed for mean spike frequency (p=0.052; **Fig. 4e**). In conclusion and in accordance with our results observed in the rodent brain, *PLCG2* downregulation impacts synaptic density and function *in vitro* in hNCs.

**Figure 4.**
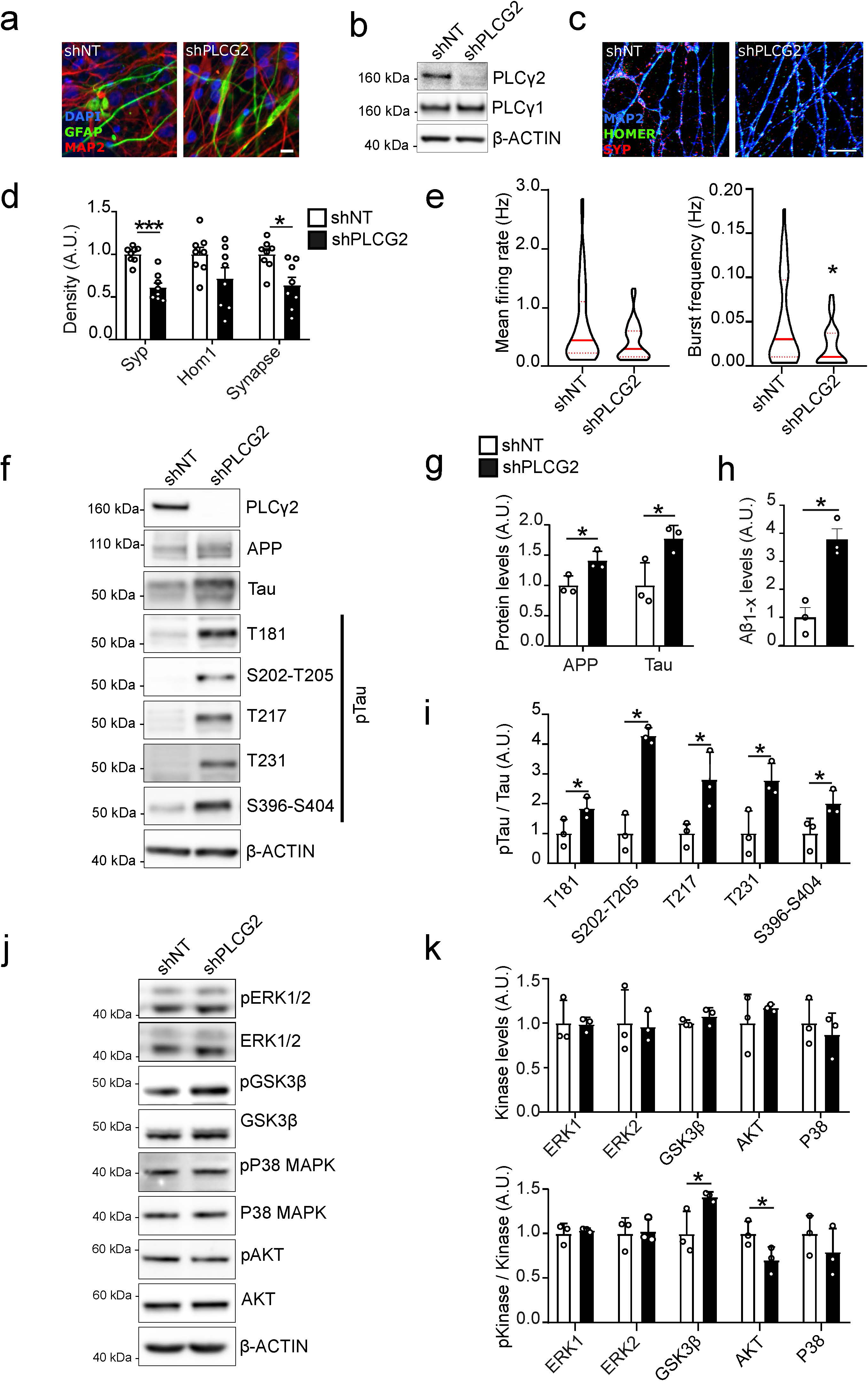
Downregulation of PLCγ2 in hNCs negatively impacts synapses and AD hallmarks. (a) Representative images of hNCs after 4 weeks of spontaneous differentiation from progenitors stably expressing either a non-targeting shRNA (shNT) or a shRNA targeting *PLCG2* (shPLCG2); scale bar = 20 µm. (b) Western blots showing the specificity of PLCγ2 downregulation *via* shPLCG2 in hNCs. (c) Immunofluorescence images of MAP2, SYP and HOMER1 synaptic markers in shNT- and shPLCG2-transduced hNCs at 4 weeks; scale bar = 20 µm. (d) Synaptic marker densities for the two conditions. Data given as mean±SEM (e) Impact of shPLCG2 on spike and burst frequencies in hNCs, measured *via* MEA. Violin plots show median and quartile lines. (f) Representative Western blots of APP, total Tau and Tau phosphorylated at T181, S202-T205, T217, T231 and S396-S404 in shNT- and shPLCG2-transduced hNCs; (g) Quantification of APP and Tau levels for the two conditions. Data given as mean ± SD, n=3. (h) Aβ peptides levels in shNT- or shPLCG2-transduced hNCs, measured by Alpha-LISA. Data given as mean±SEM, n=3. (i) Tau phosphorylation levels in hNCs expressing (or not) shPLCG2. Data given as mean±SD, n=3. (j) Representative Western blots of total and phosphorylated forms of ERK1/2, GSK3β, P38-MAPK and AKT in shNT- and shPLCG2-transduced hNCs. (k) Quantification of total and phosphorylated protein levels for these targets in shNT- and shPLCG2-transduced hNCs. Data given as mean±SD, n=3. All statistical analyses were performed using Mann-Whitney test. *p<0.05, ***p<0.001.

### Downregulation of *PLCG2* in human neuronal cultures increases both Tau phosphorylation and the levels of Aβ

We next tested the potential impact of *PLCG2* downregulation in hNCs on the different hallmarks of AD, *i.e.*, amyloid precursor protein (APP) metabolism and quantity/phosphorylation state of Tau. In shPLCG2-hNCs, a significant increase in both APP (**Fig. 4f,g**) and Aβ levels (**Fig. 4h**) was detected. *PLCG2* downregulation was also associated with a strong increase in Tau phosphorylation at multiple epitopes (**Fig. 4f,i**). Of note, an increase in total Tau protein levels was also detected (**Fig. 4f,g**). We finally investigated whether the impact of *PLCG2* downregulation on Tau hyperphosphorylation could be mediated by the regulation of ERK1/2, GSK3β, p38 MAPK or AKT kinases, known regulators of Tau phosphorylation (**Fig. 4j,k**). No difference in the total protein levels or phosphorylation statuses of ERK1/2 or p38 MAPK was observed when *PLCG2* was downregulated. However, *PLCG2* downregulation increased the level of phosphorylated GSK3β at Y216 and decreased the level of phosphorylated AKT at S473 (**Fig. 4k**). Both epitope sites are indicative of the active forms of these kinases. Altogether, these data indicate that *PLCG2* downregulation is associated with increased Aβ levels and Tau phosphorylation, the latter potentially *via* the AKT/GSK3β axis.

To confirm that the effect of *PLCG2* downregulation on AD hallmarks was specific, we performed rescue experiments by overexpressing PLCG2 in shPLCG2-hNCs. Remarkably, rescuing PLCγ2 levels almost completely restored Aβ levels, as well as total Tau and Tau phosphorylation levels (**Extended Fig. 1a-d**). Finally, the changes in GSK3β and AKT phosphorylation levels were also restored after overexpressing PLCG2 (**Extended Fig. 1a,e,f**). In conclusion, restoring PLCG2 expression in shPLCG2-hNCs can reverse the changes in components of AD hallmarks.

### Rare *PLCG2* loss-of-function variants are associated with an increase in AD risk

Importantly, our above-mentioned data suggest that the *PLCG2* downregulation may be deleterious and favor the development of AD through synaptic impairment and increased Aβ levels and Tau hyperphosphorylation. Therefore, we postulated that loss-of-function (LoF) variants responsible for a univocal reduction in *PLCG2/*PLCγ2 expression/activity might be associated with an increase in AD risk. To address this possibility, we used two independent whole exome sequencing (WES) datasets. In a Finnish sample of 527 AD cases (ADGEN) and 8,707 controls (see Material and Methods), the comparison of allele number between AD patients (2 carriers) and controls (3 carriers) indicated an overrepresentation of LoF variants in AD cases (OR = 11.0, 95% CI = 1.8-66.3; Pearson’s χ^2^ test, p=0.0009; Fisher’s exact test, p=0.03). In the ADES dataset (including 8,732 AD cases and 8,955 controls from Germany, France, Spain, the Netherlands and the United Kingdom^7^), the comparison of LoF allele number between AD patient (9 carriers) and control (1 carrier) groups also indicated an overrepresentation in AD cases (OR = 9.2, 95% CI = 1.2-72.9; Pearson’s χ^2^ test, p=0.01; Fisher’s exact test, p=0.01). Finally, a meta-analysis was performed, which, in turn, indicated that the *PLCG2* LoF variants increase the risk of AD (OR = 9.7, 95% CI = 2.0-47.7, p=0.005; **Fig. 5a** and supplementary Table S2 for a full description of the LoF variants).

**Figure 5.**
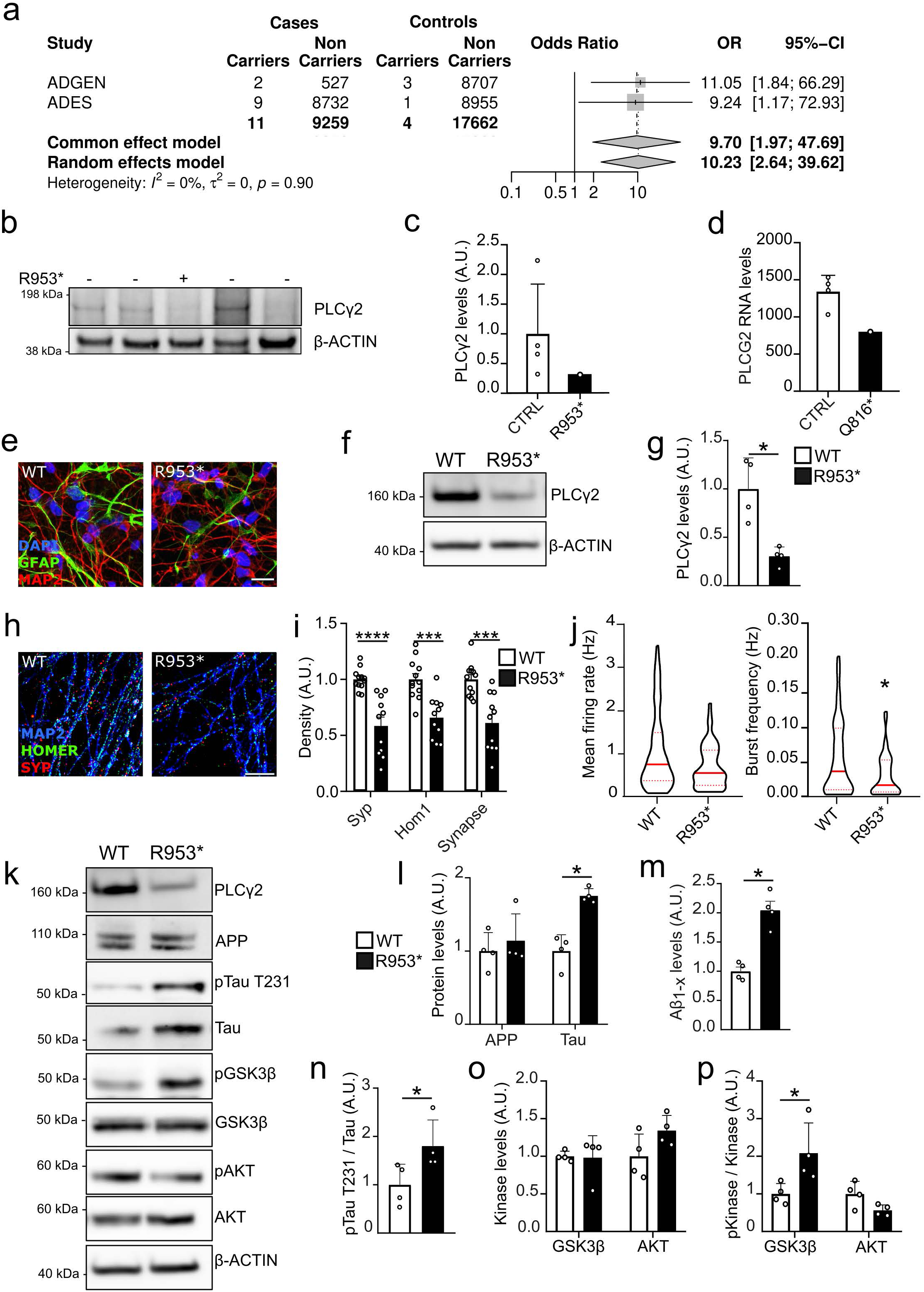
LoF nonsense variants increase AD risk and reduce *PLCG2*/PLCγ2 levels, consistently with the detrimental impact of the LoF R953* mutation on synaptic function and AD hallmarks in hNCs. (a) Meta-analysis of the ADGEN and ADES Whole Exome Sequencing data. (b) Western blot of the post-mortem brain tissue of an AD patient carrying the p.R953* variant in comparison with four other AD cases. (c) Quantification of PLCγ2 protein levels in these samples. (d) RNAseq showing 40% decrease in the blood *PLCG2* mRNA level in the p.Q816* carrier as compared to the mean of the matched control group (n=4). (e) Representative images of hNCs after 4 weeks of spontaneous differentiation from neural progenitors expressing either PLCG2^wt^ or PLCG2^R953*^; scale bar = 20 µm. (f-g) Representative Western blot (f) and quantification (g) of PLCγ2 in PLCG2^wt^ or PLCG2^R953*^ hNCs at 4 weeks. (h) Representative immunofluorescence images of MAP2, SYP and HOMER1 in PLCG2^wt^ or PLCG2^R953*^ hNCs at 3 weeks; scale bar = 10 µm. (i) Synaptic marker densities for the two conditions. Data given as mean±SEM; (j) Impact of the R953* mutation on spike and burst frequencies in hNCs, measured *via* MEA. Violin plots show median and quartile lines, n=3; (k) Representative Western blots of APP, total and T231-phospho-Tau, total and phospho-GSK3β, and total and phospho-AKT protein levels in PLCG2^wt^ and PLCG2^R953*^ hNCs. Quantification of APP and Tau protein levels (l) and relative Tau phosphorylation at T231 (m) in PLCG2^wt^ or PLCG2^R953*^ hNCs. Data given as mean±SD, n=4. (n) Aβ peptides levels in the culture medium, measured *via* Alpha-LISA and normalized by total protein levels, in PLCG2^wt^ or PLCG2^R953*^ hNCs. Data given as mean±SEM, n=4. Changes in total protein levels (o) and relative phosphorylation (p) for GSK3β and AKT in in PLCG2^wt^ or PLCG2^R953*^ hNCs. Data given as mean±SD, n=4. All statistical analyses were performed using Mann-Whitney test. *p<0.05.

### Effects of *PLCG2* LoF variants on AD pathological markers and on *PLCG2*/PLCγ2 levels

We next analyzed a post-mortem sample obtained from the temporal cortex of an AD patient carrying the *PLCG2* LoF p.R953* variant to see whether partial loss-of-function of PLCγ2 influences AD-related neuropathology. The patient who was a female was previously diagnosed as clinical AD and was classified as a Braak stage VI case according to neurofibrillary pathology in the neuropathological examination. Quantification of AD-related pathological hallmarks in the brain and CSF of this patient did not show any major alterations with respect to four Braak stage-matched AD patient samples (supplementary Table S3 and supplementary Fig. S6). Even if a large variation was observed between individuals for PLCγ2 levels in brain lysates, the PLCγ2 level was on average 70% lower in the *PLCG2* p.R953* variant carrier as compared to other AD cases (**Fig. 5b,c**). We did not detect additional specific bands for the p.R953* variant below the full-length PLCγ2. In addition, we had access to RNA extracted from the frozen blood sample of an ADGEN AD patient carrying the *PLCG2* p.Q816* variant. We randomly selected four age-matched AD patients without *PLCG2* LoF variants from the ADGEN cohort. RNA analysis of these samples revealed on average a 40% reduction in the mRNA levels of *PLCG2* in the p.Q816* variant sample as compared to the other AD samples (**Fig. 5d**). These findings collectively support the involvement of nonsense-mediated mRNA decay (NMD), as both mutations introduce premature termination codons (PTCs) that meet the established criteria for triggering this pathway^28^. However, a detailed analysis of the RNA sequencing data revealed that the Q816* T allele— corresponding to the premature stop codon—was not entirely eliminated by the NMD machinery, as it was still present in 20% of the total PLCG2 reads (data not shown). These results suggest that both LoF variants, *PLCG2* p.Q816* and *PLCG2* p.R953*, likely trigger NMD, leading to reduced mRNA and protein levels of *PLCG2*/PLCγ2.

### The *PLCG2* p.R953* LoF variant impairs synaptic function and increases both Tau phosphorylation and Aβ levels

We generated iPSCs carrying the *PLCG2* p.R953* LoF variant using CRISPR/Cas9 editing (KOLF 1.2J line, see Methods online; **Fig 5e** and supplementary Fig. S7). In *PLCG2*^R953*^-hNCs, we observed reduced PLCγ2 protein levels when compared to *PLCG2*^WT^-hNCs (**Fig 5f, g**), supporting NMD due to the p.R953* PTC. As reported in shPLCG2-hNCs, we observed that the *PLCG2* p.R953* variant associated with (i) a significant decrease in synaptic density (**Fig 5h,i**); (ii) a significant decrease in burst frequency along with a trend toward reduced mean spike frequency (p=0.061; **Fig. 5j**); (iii) a significant increase in total Tau levels and in Tau phosphorylation at T231 (**Fig. 5k-m**); (iv) a significant increase in Aβ levels in conditioned media (**Fig 5n**); and (v) a significant increase in GSK3β phosphorylation at Y216 (**Fig 5o,p**). No significant decrease in AKT phosphorylation at S473 was observed, although a similar trend was noted (–45%, p=0.057). Altogether, these data indicate that the *PLCG2* p.R953* LoF variant mirrors the phenotypic alterations previously observed following shRNA-mediated downregulation of PLCG2.

### Single nuclei (sn)-RNAseq analyses confirm that lower *PLCG2* expression affects synaptic pathways

To analyze PLCG2-dependent signaling pathways in neurons and astrocytes derived from human iPSCs independently, we conducted single-nucleus RNA sequencing (sn-RNAseq) in shPLCG2-hNCs *vs* shNT-hNCs, and in *PLCG2*^R953*^-hNCs *vs PLCG2*^WT^-hNCs. After performing quality control (supplementary Fig. S8) and clustering (**Fig. 6a** and supplementary Fig. S8), we observed a higher proportion of inhibitory neurons in shPLCG2-hNCs (**Fig. 6b**), a change mirrored in *PLCG2*^R953*^-hNCs to a lower extent. Differential expression analysis revealed a higher number of differentially expressed genes (DEGs) in shPLCG2-hNCs (**Fig. 6c**). This may be attributed to the more pronounced reduction in *PLCG2* expression as compared to the *PLCG2* R953-hNCs model, although model-specific factors are also likely to play a role, i.e., genetic background (ASE-9109 and KOLF2.1J), (ii) the mode of PLCG2 downregulation (shRNA-mediated knock-down versus an isogenic point mutation), and (iii) polyclonal versus monoclonal cell line. To disentangle the effects of reduced *PLCG2* expression from model-specific factors, we conducted separate functional enrichment analyses in each cellular model (supplementary Fig S9). Despite the models having different DEG distribution (**Fig 6c**, supplementary Tables S4 and S5), many enriched pathways overlapped, for instance those linked to synaptic function and neuronal development (**Fig. 6d, e**, and supplementary Table S6). Notably, about 25-45% of DEGs in *PLCG2*^R953*^-hNCs overlapped with those in shPLCG2-hNCs, which supports a shared PLCG2-driven signature (**Fig. 6d**). Pathway analysis of these shared DEGs revealed consistent enrichment in synaptic organization, neuronal differentiation and the calcium signaling pathway (**Fig. 6e**, and supplementary Table S6). As we observed an activation of GSK3β following *PLCG2* downregulation in our different models (**Extended fig. 1f, Fig. 4k, Fig. 5p**), we conducted regulon analyses by selecting several transcription factors that are known to depend on GSK3β activation (CREB, FOXO1, LEF1, NFATC2, NFKB, and TCF7L2)^29–33^. We found that several of these regulons were differentially represented in PLCG2-downregulated cells compared to WT cells, especially in excitatory and inhibitory neurons (supplementary Fig. S9i).

**Figure 6.**
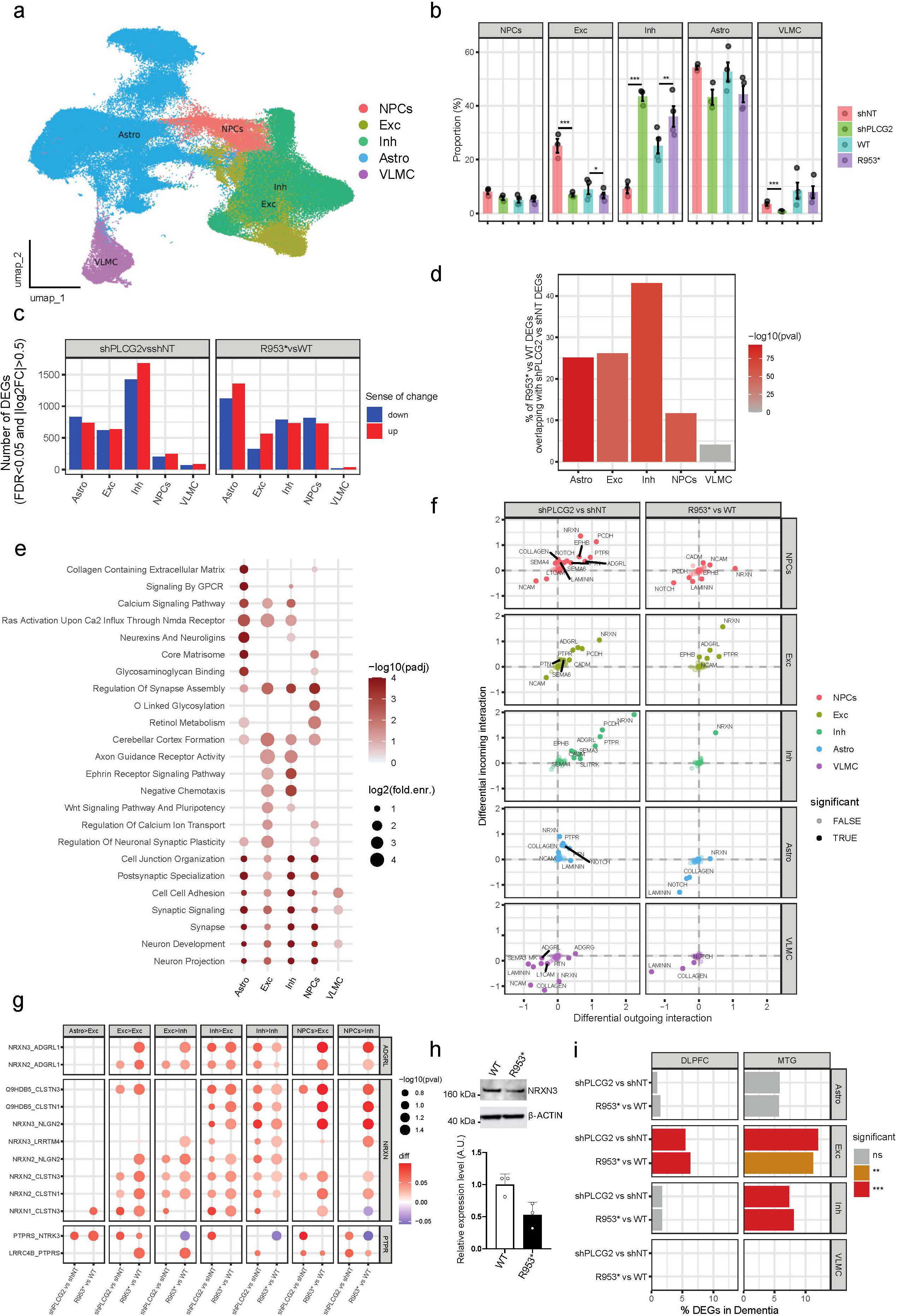
Sn-RNAseq analyses in shNT/shPLCG2 and PLCG2^wt^/PLCG2^R953*^ hNCs. (a) UMAP projection of the different cell subpopulations identified in the 140,163 transcriptomically profiled high-quality nuclei from 14 hNC cultures after QC (n=3 per group for shPLCG2 and shNT, and n=4 per group for R953* and WT). (b) Mean proportion of cell types for each condition. (*: FDR<0.05, **: FDR<0.01, ***: FDR<0.001, sccomp test^50^, adjusted for the experimental batch. (c) Number of differentially expressed genes (DEGs) comparing shPLCG2 *vs* shNT or R953* *vs* WT gene expression in each cell type. DEGs were identified according to adjusted p-value<0.05 and |log2(FoldChange)|>0.5, using pseudobulk DESeq2 analysis, adjusted for average mitochondrial RNA percentage, library batch, and ambient RNA level. (d) Common DEGs in the models, expressed as percent overlap of R953* *vs* WT DEGs with shPLCG2 *vs* shNT DEGs for each cell type. Statistical significance was assessed using the hypergeometric test. (e) Pathways enriched in the common DEGs between models. Representative pathways selected from the top100 MSigDB CP and GO gene-sets enriched (FDR<0.05, hypergeometric test) in each cell type. (f) Inferred incoming and outgoing interaction strength signaling difference comparing the two models using CellChat with CellChatDB.human database excluding Non-protein Signaling. Difference of cumulative interaction signaling received or sent by each cell type in the two models. (g) Ligand-Receptors of ADGRL, NRXN and PTPR pathways differentially interacting (p-value<0.1, Wilcoxon rank-sum test). (h) NRXN3 protein level in PLCG2^wt^ or PLCG2^R953*^ hNCs (n=3). Data given as mean±SD. Statistical analysis was performed using Mann-Whitney test, p=0.076. (i) Overlap between DEGs found when comparing PLCG2 downregulation models and DEGs reported for Dementia vs non-Dementia groups in the SEA-AD postmortem brain snRNA-seq in two different brain regions. DLPFC: dorsolateral prefrontal cortex; MTG: middle temporal gyrus. Over-representation analysis using hypergeometric test using as background all genes tested for differential expression in the PLCG2 models. Astro, astrocytes; Exc, excitatory neurons; Fib, fibroblasts; Inh, inhibitor neurons; NPCs, neuroprogenitor cells; VLMC, vascular leptomeningeal cells.

We also investigated intercellular communication using CellChat^34^. On average, 727 ligand–receptor (L/R) interactions were inferred *per* sample, grouped into 89 signaling pathways, with excitatory– inhibitory neuron interaction being strongest (supplementary Fig. S10b). Differential analysis revealed 18 and 10 altered L/R signaling pathways in shPLCG2-hNCs *vs* shNT-hNCs and *PLCG2*^R953*^-hNCs *vs PLCG2*^WT^-hNCs, respectively, with three main pathways depending on NRXN, ADGRL and PTPR, upregulated or downregulated in both models in neurons (**Fig. 6f,g** and supplementary Fig. S10). Among them, NRXN-dependent pathways were strongly upregulated in excitatory and inhibitory neurons across models (**Fig. 6g**). To strengthen the potential implication of neurexins, we selected NRXN3 for further validation, given that this isoform has been most consistently implicated in AD pathogenesis^35–39^. We observed a trend towards lower levels (–47%, p=0.076) of NRXN3 protein in PLCG2^R953*-^hNCs compared to PLCG2^WT^-hNCs (**Fig. 6h**), although NRXN3 mRNA levels were not different in neurons from PLCG2^R953*^-hNCs compared to PLCG2^WT^-hNCs (supplementary Table S4). This supports that NRXN3-dependent L/R signaling pathways are altered in our cellular models. To note, we also observed Laminin signaling downregulated in vascular leptomeningeal cells (VLMCs)–astrocytes communication, although less consistently (supplementary Fig S10c).

Finally, we sought to detect a potential AD-related transcriptomic signature in by comparing our sn-RNAseq results with publicly available snRNAseq databases and found a significant overlap in DEGs with the SEA-AD snRNAseq dataset^40^, predominantly in excitatory neurons and, to a lesser extent, in inhibitory neurons (**Fig 6i**).

Altogether, these results suggest that *PLCG2* downregulation perturbs neuronal and synaptic functions, at least partly *via* neuron-neuron communication, with NRXN/ADGRL signaling emerging as a key candidate pathway. In addition, these observations suggest that PLCG2 downregulation can participate to generate, in part, an AD-related transcriptomic signature in excitatory neurons.

## DISCUSSION

To determine if and how AD genetic risk factors contribute to synaptic failure, we developed an HCS to systematically characterize the impact of the downregulation of GWAS-defined genes on synaptic density. Among the genes with the strongest impact on synaptic density, *PLCG2* was one of the best hits (despite the absence of microglia in our HCS). In AD, *PLCG2* has been mostly studied in microglia and shown to be implicated in multiple microglia-related signaling pathways^18–21^. However, strong evidence supports a role for PLCγ2 in synaptic function: (i) we and others found that *PLCG2* is also expressed in neurons^22–24;^ (ii) we observed that Plcγ2 is localized to both the pre- and post-synaptic compartments; (iii) we demonstrated that PLCγ2 downregulation impacts synaptic density and neuronal electrophysiological properties using four distinct experimental models (and different shRNAs).

To further support the importance of PLCγ2 and its downregulation in AD pathophysiology, we determined that very rare LoF variants in *PLCG2* are associated with an increased risk of AD in two independent WES datasets. Two of these LoF variants, p.Q816* and p.R953* were associated with a decrease in *PLCG2*/PLCγ2 expression in the blood or brain and we confirmed such a decrease in PLCγ2 expression in *PLCG2*^R953*^ hNCs. Remarkably, we also observed that the R953* variant leads to increased Tau phosphorylation and Aβ levels, suggesting that PLCγ2 participates in the development of the two main hallmarks of AD. The shRNA-mediated downregulation of *PLCG2* also has the same impact as the variant, supporting the idea that a decrease in *PLCG2* activity is harmful. This is consistent with the fact that the functionally hypermorphic *PLCG2* p.P522R variant is protective against AD^24^. Interestingly, a recent study using the cross-bred 5xFAD; Plcγ2-P522R knock-in mice revealed that the hypermorphic P522R variant improves impaired synaptic function as well as behavioral deficits observed in 5xFAD mice^21^. Collectively, these observations suggest mechanisms where the sustained activity of PLCγ2 becomes instrumental when the risk of AD (protective *vs* deleterious) is considered not only in microglia, but also in neurons.

Characterizing the potential mechanisms through which PLCγ2 influences key hallmarks of AD in hNCs is of particular interest. We first focused on kinases known to be regulators of Tau phosphorylation. Interestingly, our results suggest that GSK3β may play a role, among other factors, in mediating the cellular response downstream of PLCγ2 downregulation. GSK3β is already known to be involved in the regulation of APP metabolism, Tau phosphorylation and synaptic functions^41–43^, making it a potential link between PLCγ2 and the different phenotypes associated with its downregulation. Even though we did not demonstrate that GSK3β is a direct intermediate of the PLCγ2 impact on AD hallmarks, activation of GSK3β was fully abolished when PLCγ2 level was restored, supporting a specific impact of PLCγ2 on this phenotype. Of note, neurons derived from iPSCs obtained from sporadic AD cases also exhibit elevated Tau hyperphosphorylation, increased Aβ levels, and GSK3β activation^44^. AKT emerged as a key target, as its membrane recruitment and activation depend on PLCγ2-regulated substrate diacylglycerol (DAG). DAG-driven PKC activation can enhance AKT signaling and loss of PLCγ2 function may thus reduce AKT activity, as observed. However, in the brains of Plcγ2-P522R knock-in mice, Akt1/2 phosphorylation was reduced, casting doubt on this hypothesis^18^.

We next performed snRNA-seq on both hNC models and found that numerous pathways related to synaptic and neuronal functions were commonly affected. This is consistent with the observed reductions in synaptic density and function following *PLCG2*/PLCγ2 downregulation. Notably, the geneset “regulation of neuronal synaptic plasticity” was preferentially altered in excitatory neurons. In addition, multiple pathways were impacted across different cell types, suggesting complex intercellular interactions within hNC cultures. To investigate whether *PLCG2*/PLCγ2 downregulation preferentially disrupts interactions between specific cell populations, we performed a cell-cell communication analysis. This revealed that ADGRL and NRXN signaling were preferentially affected within neurons across both models. Both pathways are involved in synaptic function and psychiatric disorders^45–47^. NRXN signaling is of particular interest, as neurexins have been shown to regulate APP synaptic functions and/or modulate Aβ-induced synaptotoxicity^36,37,48^.

Importantly, there are some limitations related to our cellular models that need to be mentioned. Firstly, *PLCG2* downregulation was associated with accelerated neuronal maturation in all models (see supplementary Fig. S9e). We thus cannot exclude that this difference in maturation impacts the observed modifications in phenotypes following *PLCG2* downregulation. However, it is worth noting that the downregulation of *Plcg2 in vivo*, which leads to profound changes in DG granule cell morphology and physiology, was performed in adult (>3 months old) mice, thus precluding a potential developmental effect that could be argued to explain part of the synaptic phenotype observed in rat PNCs and hNCs. Secondly, although the CellChat analyses and observation of enriched AD-related transcriptomic signatures in neurons both support the idea that *PLCG2* downregulation in neurons is responsible for the observed phenotypes, we cannot rule out the possibility that *PLCG2* downregulation in astrocytes contributes to them.

In conclusion, our results reveal a novel role for the AD genetic risk factor *PLCG2*. Downregulation of *PLCG2* specifically in neurons results in a signaling cascade that leads to the activation of GSK3β. This may promote increased Aβ secretion, Tau hyperphosphorylation, and synaptic dysfunction. These results are consistent with previous reports suggesting that PLCγ2 may modulate Tau pathology and disease progression in patients with mild cognitive impairment^49^. As an emerging major genetic risk factor for AD and other neurodegenerative diseases, *PLCG2* represents a promising therapeutic target. Future studies will be critical to delineate PLCγ2-dependent pathways in both microglia and neurons in order to identify relevant druggable targets.

## Acknowledgement

The authors thank the BICeL platform of the Institut Biologie de Lille and the Vect’viral platform (INSERM US 005 – CNRS 3427 – TBMCore, Université de Bordeaux, France). The authors thank Karine Blary at the IEMN Lille for the microfabrication work. The authors also thank Sisko Juutinen, Eija Rahunen, and Minna Turunen for excellent technical support with immunohistochemistry. Fluorescent microscopy of human brain samples was carried out with the support of UEF Cell and Tissue Imaging Unit, University of Eastern Finland, Finland. This work has been funded by JPND (JPND2019-466-128, project PGM-AD; JPND2023-1822-030, project AD-PLCG2), ANR (GENSYNALZ, ANR-21-CE16-0010; AD-PLCG2, ANR-24-CE14-5376-01), COEN (grant number 6005), Alzheimer (grant number 903716), Fondation Claude Pompidou, Fondation pour la Recherche Médicale (ALZ201912009628), Fondation Recherche Alzheimer, Fondation Vaincre Alzheimer (grant number FR-22032), and the French RENATECH network (P-18-02737). This work was also supported by Lille Métropole Communauté Urbaine and the French government’s LABEX DISTALZ program (Development of innovative strategies for a transdisciplinary approach to Alzheimer’s disease), European Union under the European Regional Development Fund (ERDF), Hauts de France Regional Council (contract469 no.18006176), MEL (contract_18006176), and the French State (contract no. 2018-3-470 CTRL_IPL_Phase2). This work has been funded by Research Council of Finland (grant number 330178, 338182, 355604, 334802), the Sigrid Jusélius Foundation, Jane and Aatos Erkko Foundation, the Strategic Neuroscience Funding of the University of Eastern Finland, Faculty of Health Sciences of University of Eastern Finland and Alzheimer’s Association (ADSF-24-1284326-C and AARG-22-926152). This work was also funded by ERC-StG Metabo3DC (101042759) to JTH et LabEx EGID (ANR 10-LABX-0046). This work was also supported by the National Institute of Health (R01AG082362, R01AG083941), Bright Focus Foundation, The Edward N. & Della L. Thome Memorial Foundation Awards Program in AD Research, Health Resources in Action. EG acknowledges support by the Luxembourg National Research Fund (INTER/JPND23/17999421/AD-PLCG2) and by the Luxembourg Fondation Wivine. This work is supported by the UK Dementia Research Institute [award number UKDRI-Car001] through UK DRI Ltd, principally funded by the Medical Research Council.

## Author’s Contribution

**Conceptualization**: J.C., J.D., J-C.L; **High content screening**: A.C., T.M., D.K., J.C., J-C.L.; **Cell culture experiments**: A.C., F.R., N.L-W., D.S-W., A.P., M.L., F.L., K.G., P.W., A.R.M.F., C.N., A. F-R., C.G., E.L., O.S., N.M., M.R.C., A-M.A, F.D., D.K., J.D.; **iPSCs QC**: N.L-W., B.G-B., M.M., J.D.; **Electrophysiological analyses**: A.R., A.E., S.D., C.M.; **Brain, blood and CSF analyses in human samples**: M.T., H.M., R.M.W., H.J., P.M., S.H., T.R., H.S., M.Hil.; **RNAseq analyses:** A.P., A.G., V.G., L-I.C., V.G., J.TCW., E.G., D.E., J.T.H; **WES sequencing data**: M.T., H.M., M.Hul., P.A., C.C., O.D-I., H.J., T.K., M.K., J.H., S.H., H.H., P.M., G.N., S.M., M.W., A.R., T.R., A.P., R.S., H.S., J.V.S., J.W., R.M.W., M.Hil., J-C.L.; **Genetic Analysis**: M.Hul., T.K., S.H., B.G-B., M.Hil., J-C.L. **Writing**: A.C., F.R., M.T., A.R., A.P., H.M., D.K., C.M., J.C., M.Hil., J.D., J-C.L.

## Availability of data and material

Cellular models used and/or analyzed in the current study are available from the corresponding authors upon reasonable request. snRNA-seq and WGS datasets will be made available before publication.

## METHODS ONLINE

### Primary neuronal culture

Primary hippocampal and cortical neurons were obtained from P0 Wistar rats, according to previously described procedures with minor modifications^51^. Culture media and supplements were from Gibco, ThermoFisher Scientific, unless mentioned otherwise. Briefly, hippocampi and cortex were isolated, washed with ice-cold dissection medium (HBSS supplemented with sodium pyruvate, HEPES, and penicillin/streptomycin), and trypsinized (2.5%) 10 min at 37°C. Trypsin was then inactivated with dissociation medium (MEM supplemented with inactivated FBS, Glutamax, D-glucose (Sigma), MEM vitamins, and penicillin/streptomycin) containing DNase (5 mg/ml, Sigma). Tissues were then dissociated in culture medium (Neurobasal A supplemented with glutamax and B27) to obtain a homogenous cell suspension, followed by centrifugation (200× g for 8 min). Cells were resuspended in the culture medium and seeded into plates that were pre-coated with poly-D-lysine (0.1 mg/mL, Sigma) in a borate buffer solution (0.31% boric acid and 0.475% sodium tetraborate, pH = 8.5). Cells were resuspended in culture medium and seeded in plates precoated with poly-D-lysine (0.1 mg/mL, Sigma) in borate buffer (0.31% boric acid, 0.475% sodium tetraborate, pH = 8.5) at a density of 50,000 cells/cm^2^ for 384-well plates (Greiner bio-one), 100,000 cells/cm^2^ for 24-well plates, 125,000 cells/cm^2^ for 10 cm culture dishes or at 800,000 cells/cm^2^ for microfluidics. For MEA 96-well plates, a 5 µL drop containing 15,000 cells was placed on the top of the electrodes, and medium was added after a minimum of 5 minutes to allow the cells to attach. For all types of culture, media was fully replaced after 24 h and cells were maintained in a tissue culture incubator at 37°C with 5% CO2 for 21 days.

### Generation and quality control of the PLCG2 p.R953* KOLF2.1J hiPSC line

Isogenic hiPSCs heterozygous for the p.R953* variant (R953*) were generated *via* CRISPR-Cas9 editing of the KOLF2.1J cell line^52^ (wild-type, WT) by Applied StemCell (CA, USA). The guide RNA sequence was GAGCGGATTTCTCGGAAGTCAGG and the single-stranded oligodeoxyribonucleotide (ssODN) repair template sequence was GAAGTACTGATGACCTTTTTCTCTGTGTGCAGAAAATCATGACTTCTGAGAAATCCGCTCCTTTGTGGAGACGAA GGCTGACAGCATCATCAGACAGAAGCCCGTCGACCTCCTGAAGTACAATCAAAAGGGCCTGACCCGCGTCTACCCGAAGGGACAAAGAGTTGACTCTTCAAACTACGAGGGGTTCCGCCTCTGG. The successful introduction of the intended C>T heterozygous substitution (R953*) was verified using Sanger sequencing with the following primers: GTGGGAAAACTGGCGCTCCT and ACCATCTGGAACGCGACAG. The typical morphology of the CRISPR-edited iPSCs and the presence of pluripotency factors (e.g., OCT4 and SSEA4) were verified using live brightfield microscopy and immunofluorescence staining, respectively. Genomic DNA (gDNA) was extracted from the CRISPR-edited iPSCs and the parental KOLF2.1J iPSCs using NucleoSpin Tissue (Macherey-Nagel) according to the manufacturer’s instructions. Fingerprinting of the cell lines was performed to rule out any mix-ups during the editing process or contamination by other cell lines. This was performed by assaying a microvariation at the human D1S80 locus on chromosome 1^53^. Allele variability between individuals was visualized by analyzing PCR products (primers: GTCTTGTTGGAGATGCACGTGCCCCTTGC and GAAACTGGCCTCCAAACACTGCCCGCCG) on 2% agarose gel. Potential off-target and on-target genomic alterations introduced by CRISPR editing were assessed using quantitative genomic PCR (qgPCR) as described previously^54^ and whole-genome sequencing (WGS). qgPCR employed the following primers and probe: forward primer GTGGGAAAACTAGGCCCCTT, reverse primer ACCATCTGAGAACCGCACAG, and the fluorescently labeled probe 5’ 6-FAM/CCGAGAAAT/ZEN/CCGCTCCTTTGTGGA/IBFQ 3’ (PrimeTime Eco Probes, IDT), with Human TERT on chromosome 5 serving as a reference gene (Human TaqMan Copy Number Reference Assays, Thermo Fisher Scientific, cat. no. 4403316). Karyotypic integrity was confirmed by qPCR using the hPSC Genetic Analysis kit (StemCell Technologies, cat. no. 07550) and WGS.

WGS of parental and edited KOLF2.1J lines was outsourced at Beijing Genomics Institute (BGI). Reads were processed using the nf-core sarek pipeline (v3.5.1)^55^, incorporating fastp (v0.24.0) for quality control, dragmap (v1.2.1) for alignment to the reference genome GRCh38, and Picard MarkDuplicates (v3.3.0) for duplicate marking. Variant calling was conducted with GATK HaplotypeCaller (v4.6.1.0), chromosome copy number was estimated via tiddit (v6.6.1), and variants were annotated with Ensembl’s VEP (v111). Structural variant (SV) detection was conducted using three software: DELLY (v1.7.2)^56^, Manta (v1.6.0)^57^, and Smoove (v0.2.8; https://github.com/brentp/smoove) coupled with duphold (v0.2.3)^58^. Alignment of reads was performed by sarek using BWA (v0.7.18) instead of dragmap as these three SV callers perform better with this aligner to detect discordant and split reads. Joint calling was performed for each software using default parameters including both parental and edited KOLF2.1J lines. Quality control was performed and SVs were retained only if they met the following criteria: a quality score (QUAL) above 200, read coverage below 100, a minimum of five supporting reads per allele and an allele balance between 0.2 and 0.8 for heterozygous calls. For homozygous calls, at least ten supporting reads and an allele balance of 0.9 or higher were required. Software-specific quality thresholds were also applied in DELLY (FILTER=PASS, excluding LowQual genotypes), Manta (FILTER=PASS, genotypes classified as PASS or HomRef), and Smoove (GQ>20, SHQ=4 for heterozygous calls, deletions with DHFFC<0.7, and duplications with DHFFC>1.25). Resulting SVs from each tool were then merged using SURVIVOR (v1.0.7)^59^, with a maximum allowed breakpoint distance of 500 base pairs and considering both SV type and strand orientation. The final set of SVs was restricted to those identified by all three tools (or by both DELLY and Manta in the case of insertions, as Smoove does not detect this SV type) and only those whose genotype assignments between software were consistent across software (i.e. heterozygous versus homozygous and non-missing) were retained. Sample contamination was evaluated using VerifyBamID (v2.0.1). Default parameters were applied for all pipeline steps.

WGS analyses confirmed the precise introduction of the intended C>T heterozygous substitution (R953*) at chromosome 16, position 81,936,183 (GRCh38). Five additional single nucleotide variants were detected within the *PLCG2* gene region (chr16:81,738,248–81,962,685). Two of them (chr16:81,936,284:C>T and chr16:81,936,287:A>G) were homozygous and intentionally introduced as synonymous mutations in the design to disrupt Cas9 target sites and prevent re-cutting post-ssODN repair. The three other mutations (chr16:81,936,175:C>A, chr16:81,936,218 C>T, and chr16:81,919,070 A>T) were heterozygous mutations. The chr16:81,936,175:C>A is a missense variant (P950H) but this mutation is in phase with the intended C>T heterozygous substitution we introduced (R953*); its impact is therefore negligible. The chr16:81,936,218 C>T and chr16:81,919,070 A>T mutations are unlikely to affect protein function since the first one is a synonymous variant and the second one is located in an intron. Quality control confirmed normal ploidy and the absence of CRISPR-induced large deletions, insertions, chromosomal rearrangements, or loss of heterozygosity. Furthermore, analysis of the 33 off-target sites predicted by CRISPOR^60^ showed that no off-target single nucleotide or structural variants introduced by CRISPR-Cas9 editing. An overview of the QC performed on CRISPR-edited iPSCs is provided in supplementary Fig. S7.

### Human iPSC-derived neurons and astrocytes co-culture

Culture media and supplements were obtained from Stem Cell Technologies, unless mentioned otherwise. Available ASE-9109 hiPSC cell line and CRISPR-edited heterozygous PLCG2 p.R953* KOLF2.1J hiPSC cell line (Applied Stem Cell Inc. CA, USA) were maintained in non-treated cell culture dishes/plates pre-coated with 10 µg/ml of vitronectin fresh-diluted in Cell Adhere Dilution Buffer, in mTeSR+ medium fully changed daily. The ASE-9109 and KOLF2.1J NPCs were generated using the embryoid body and monolayer protocols from Stem Cell Technologies, respectively. In both protocols, iPSCs were induced into neural stem cells through dual SMAD inhibition, where progenitors upregulate *PAX6* and *EMX1* genes, suggesting a forebrain-like phenotype^61^. NPCs were maintained in complete Neural Progenitor Medium (NPM, supplemented with supplements A and B), in treated cell culture dishes pre-coated with 0.001% poly-L-ornithine (PLO) diluted in water and 10 μg/mL laminin diluted in 0.01 M PBS with Ca^2+^ and Mg^2+^. Full media changes were performed daily. To generate NPCs stably expressing shRNAs non-targeting or targeting PLCG2 gene, NPCs derived from ASE-9109 hiPSCs were transduced with lentiviral shRNAs carrying an antibiotic resistance cassette (pLV-Hygro-U6>non-targeting_shRNA#1: CCTAAGGTTAAGTCGCCCTCG and pLV-Hygro U6>shPLCG2_shRNA#1: CAGTTCCATCTCTTCTATAAA, Vector Builder) 4 h after seeding at Multiplicity of Infection (MOI) 4. Two days after the transduction, Hygromycin (500 µg/mL, Invitrogen) was added to the cell medium and kept until selection was complete.

Mixed cultures of human induced neurons and astrocytes were obtained from shRNA-transduced NPCs after 3/4 weeks of spontaneous differentiation. For immunoblotting and immunostaining, 70,000-100,000 NPCs/wells were plated in PLO/Laminin pre-coated 24-well cell imaging plates (Eppendorf, 0030741005) or 24-well cell culture plates (Greiner, 662160). For microfluidic devices, 30,000-40,000 NPCs for synaptic density analysis were seeded in both chambers of pre-coated PLL/Laminin microfluidic chips. For MEA plates, a 5 µL drop of 25,000-30,000 cells was placed on the top of the electrodes, and medium was added after a minimum of 30 minutes to allow the cells to attach. Cells were kept in complete NPM containing 10 μM of Y-27632 ROCK Inhibitor for 24 h, before adding an equal volume of complete BrainPhys medium (supplemented with N2, SM1, BDNF, GDNF, laminin, dibutyryl-cAMP, and ascorbic acid)^62^. Cells were maintained in a tissue culture incubator at 37°C with 5% CO_2_, and half-media changes were performed bi-weekly with Brainphys for 4 weeks. Our snRNA-seq data confirmed that these cultures showed a predominant forebrain identity, as well as a possible differentiation towards a hippocampal granule lineage (supplementary Fig. S8d).

### HCS shRNA library

A HCS was performed to assess the impact of a lentiviral Mission pLKO,1-shRNA library targeting 198 AD-associated genes on synaptic density. The detailed methodology of our HCS was published elsewhere^16^. The screening was organized in two parts: (i) 105 genes reported in 2019^63^, and (ii) 93 genes reported in 2022^6^. These genes were selected using The Allen Brain Human MTG SmartSeq Dataset (https://portal.brain-map.org/atlases-and-data/rnaseq) to limit our analysis to genes expressed in brain cells. Non-coding genes, and genes with no rat homologs were also excluded. PNCs were transduced at MOI 2 or 4, and the HCS was performed three times using three independent neuronal cultures. A shRNA targeting Synaptophysin (shSyp, Mission, NM_009305 TRCN0000379864, GGTGGTTATCAACCCGATTAC), and a non-targeting shRNA (shNT, Mission, shC002V, CAACAAGATGAAGAGCACCAA) were used as positive and negative controls, respectively.

### shRNA and cDNA transduction of neuronal cultures

Primary neuronal cultures (PNCs) were transduced on DIV1 with lentiviral Mission pLKO,1-shRNA vectors (Merck) at MOI2 or MOI4. Viral constructs were added to pre-warmed supplemented culture media with 4 μg/mL polybrene (hexadimethrine bromide, Sigma). Culture media was removed and transduction mixture was added to each well to reach 20 µL final volume for 384-well plates or 250 μl final volume for 24-well plates. PNCs were incubated for 6 h before equal volumes of fresh pre-warmed culture media were added.

For Neural Progenitor Cells (NPCs) transduction with shNT and shPLCG2, viral constructs were diluted in cell culture medium and directly added to NPCs. For rescue experiments, shNT NPCs and shPLCG2 NPCs were transduced on the first day of the differentiation with a lentiviral control vector (pLV[Exp]-CAG>{empty}/Myc:T2A:EGFP, Vector Builder) or a lentiviral vector expressing a shRNA-resistant PLCG2 cDNA (pLV[Exp]-CAG>{PLCG2)}/Myc:T2A:EGFP, Vector Builder).

### Immunostaining

Cells were fixed with 4% paraformaldehyde (EMS, Hatfield, PA) for 20 min at RT, permeabilized with 0.3% Triton X-100 in PBS for 5 min at RT, and blocked with 5% normal donkey serum (Jackson ImmunoResearch) and 0.1% Triton X-100 for 1 h at RT. Alternatively, neurons in 384-well plates were blocked with 2.5% BSA and 0.1% Triton X-100 for 2 h at RT. Cells were washed with PBS between each step. For 384-well plates, washes were automated using an automated platform (Works automation control software (Agilent), Direct Drive Robot (Agilent), Plate washer (BioTek, EL406)). After blocking, neurons were incubated overnight at 4°C with the following primary antibodies: Chicken or Guinea-Pig anti-Homer 1 (Synaptic Systems, 160006 and 160004), Guinea-Pig or Mouse anti-Synaptophysin (Synaptic Systems, 101004 and 101011), Mouse or Chicken or Guinea pig anti-MAP2 (Synaptic Systems, 188011, 188006 and 188004), Rabbit anti-alpha Tubulin 4 (Abcam, ab18251), Guinea-Pig anti-GFAP (Synaptic Systems, 173004). After washes with PBS, cells were incubated 2h at RT with the following secondary antibodies: Donkey anti-Chicken Alexa 488 or 647 (Jackson Immunoresearch, 703-545-155 and 703-605-155), Donkey anti-Rabbit Alexa 488 or 647 (Jackson Immunoresearch, 711-645-152 and 711-605-152), Goat anti-Guinea-Pig Alexa 555 (Life technology, A21435), Donkey anti-Guinea-Pig Alexa 405 or 594 (Jackson Immunoresearch, 706-475-148 and 706-585-148), Donkey anti-Mouse Alexa 488 or 594 or 647 (Jackson Immunoresearch, 715-545-151, 711585-151 and 715-605-151).

For immunostaining of hippocampal sections from animals injected with lentiviral vectors. At the terminal timepoint mice were transcardially perfused with ice-cold saline and 4% paraformaldehyde under lethal dose of ketamine-xylazine anesthesia. Then, brains were isolated and post fixed in 4% paraformaldehyde and 40 µm thick coronal sections of hippocampus were prepared in vibratome (VT-2000, Leica) which were then incubated overnight at 4°C with following primary antibodies: Mouse anti-GFP (ab290, abcam), Mouse anti-PLCγ2 (sc-5283, Santa Cruz), Rabbit anti-iba1 (234 008, Synaptic System), Mouse anti-NeuN (1B17, abcam). Further, slices were incubated with secondary antibodies following 3 washes (5 min each) with PBS at RT for 2 h. If needed slices were counterstained with DAPI for nuclear localization. Finally, stained sections were imaged using a Leica DM5500 TCS SPE confocal microscope.

### HCS imaging and image analyses

384-well plates were imaged using IN Cell Analyzer 6000 Cell Imaging System (GE Healthcare) equipped with a Nikon 60× 0.95 NA objective and a CMOS camera. 16 images (2048 × 2048 pixels) per well were acquired in 3 stacks (z-stack interval: 0.5 µm) from 3 channels (dsRed, FITC, and Cy5). Customized image analysis software (Columbus; PerkinElmer) was used for the image analysis. MAP2 staining was used to define network area and Synaptophysin (Syp) and Homer1 spots were detected within this area. A custom MATLAB (MathWorks, Natick, MA) process was used to exclude outlier fields in terms of network area or staining intensity, and to assign each Homer1 spots to the nearest Syp spot within a 1 µm cut-off distance. The density of Syp spots assigned by at least one Homer1 spot was used to calculate the synaptic density. For each well, data was normalized by the mean of non-transduced wells from the same plate. Wells with shRNA-induced toxicity, as defined by normalized network area < 0.4, were excluded from the analysis. Assay quality was assessed using β-score calculated according to 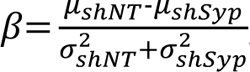 where μ and σ are the mean and standard deviation of Syp density in control wells treated with shNT and shSyp, and only plates with a β-score > 1 were analyzed. After all screenings were performed, mean and standard error of the mean (SEM) of normalized synaptic density were calculated for each shRNA that did not affect network quality in at least 2 screenings. shRNAs that potentially affect synaptic density were determined as those belonging to the top or bottom 2.5% tiers.

### Microfluidic chip fabrication

Masters of microfluidic devices were fabricated through photolithography as previously described^64^. Devices were cast from polydimethylsiloxane (PDMS, Sylgard 184, Dow Corning, Midland, MI) and irreversibly bonded to glass coverslips using an O_2_ plasma generator (Diener, Ebhausen, Germany). Prior to cell culture, devices were sterilized under UV (Light Progress, Anghiari, Italy) for 20 min, and coated either with 0.1 mg/ml poly-D-lysine (PDL, Sigma) diluted in borate buffer for PNCs or with poly-L-lysine (PLL, Sigma) diluted in borate buffer followed by 10 µg/mL laminin (CC095, Sigma-Aldrich) diluted in 0.01 M PBS with Ca^2+^ and Mg^2+^ (Gibco, ThermoFisher Scientific) for NPCs. For quantification of synapses, tricompartment microfluidic devices that isolate the neuronal network from cell bodies were used^17^. At least two devices were employed for each experimental condition and at least three independent cultures were performed.

### Synapse imaging and analysis

Synaptic compartments of microfluidic chips were imaged *via* Spinning Disk XLight V3 microscope (Nikon), using a 60× objective and a *z*-stack interval of 0.2 µm. Images were deconvoluted using Huygens Professional (Scientific Volume Imaging) and analyzed using Imaris (Bitplane, Zürich, Switzerland) by reconstructing Syp and Homer1 puncta in 3D. The relative positions of all puncta were analyzed using the custom MATLAB process described before to evaluate the synaptic density^17^. Statistically significant outliers were calculated and excluded, using the median absolute deviation method (±3 MAD). Immunostained 24-well plates were imaged *via* Spinning Disk XLight V3 microscope (Nikon), using a 40× objective and a *z*-stack interval of 0.3 µm.

### Synapse fractionation

7×10^7^ primary rat cortical cells were resuspended in a cold buffer containing 0.32 M sucrose and 10 mM HEPES, pH = 7.4, and were centrifuged at 1000× *g* for 10 min. The resulting supernatant was concentrated at 12,000× *g* for 20 min to remove the cytosolic fraction. Then, the resulting pellet was resuspended in a second solution (4 mM HEPES, 1 mM EDTA, pH = 7.4) and centrifuged twice at 12,000× *g* for 20 min. The new pellet was incubated in a third solution (20 mM HEPES, 100 mM NaCl, 0.5% Triton X-100, pH = 7.2) for 1 h at 4°C with mild agitation and then centrifuged at 12,000× *g* for 20 min. The supernatant collected corresponds to the triton-soluble non-PSD fraction. The remaining pellet was resuspended and incubated in a buffer containing 20 mM HEPES, 0.15 M NaCl, 1% Triton X-100, 1% deoxycholic acid, 1% SDS, pH = 7.5 for 1 h at 4 °C, and then centrifuged at 10,000× *g* for 15 min to obtain a supernatant containing the triton-insoluble PSD fraction. The cytosolic, non-PSD, and PSD fractions were then analyzed by immunoblotting.

### Immunoblotting

Protein quantification was performed using the BCA protein assay (ThermoFisher Scientific). Equal amounts (5–20 μg) of cell lysate were collected in RIPA buffer (0.05 M Tris, 0.15 M NaCl, 1% NP-40, 0.1% SDS, 1 mM sodium orthovanadate, and 0.5% sodium deoxycholate, pH = 7.4) containing protease inhibitors (Complete mini, Roche Applied Science, Penzberg, Germany), lithium dodecyl sulfate (ThermoFischer Scientific), and reducing agent (Invitrogen). Samples were sonicated at 60% and denaturated at 95°C for 5 min. Protein extracts were separated in 4-12% Bis-Tris, or 3-8 % Tris-Acetate SDS–polyacrylamide gels (NuPAGE, Thermo Scientific) and transferred on to nitrocellulose membranes (Bio-Rad). Membranes were blocked in SuperBlock (ThermoFisher Scientific) for 1 h at RT and incubated with primary antibodies at 4°C overnight. The following day, membranes were incubated with HRP-conjugated secondary antibodies for 2 h at RT and revealed using Immobilon Classico Western HRP Substrat (Millipore) and the Amersham Imager 600 (GE Life Sciences). Optical densities of bands were quantified using ImageJ. The following primary antibodies were used: Chicken anti-Synaptophysin (Synaptic System, 101006), Guinea-Pig anti-PSD95 (Synaptic System 124014), Chicken anti-GAPDH (Millipore, AB2302), Rabbit anti-USP6NL (ThermoFisher Scientific, PA575944), Mouse anti-PSMC3 (ThermoFisher Scientific, MA538612), Rabbit anti-CSNK1G1 (ThermoFisher Scientific, PA5-205268), Rabbit anti-CYB561 (Merck, SAB2100516), Rabbit anti-SNX1 (Abcam, ab134216), Rabbit anti-ICA1L (ThermoFisher Scientific, 16828054), Mouse anti-OPLAH (Santa-Cruz, sc-398205), Mouse anti-NCK2 (Santa-Cruz, sc-20020), Mouse anti-PLCG2 (Santa-Cruz, sc-5283), Rabbit anti-PLCG1 (Abcam, ab76155), Rabbit anti-APP (Sigma, A8717), Rabbit anti-hTau (Dako, A0024), Mouse anti-pTauS396/404 (P. Davies, PHF1), Mouse anti-pTauT231 (P. Davies, RZ3), Mouse anti-pTauS202/T205 (ThermoFisher Scientific, MN1020), Rabbit anti-pTauT217 (ThermoFisher Scientific, 10536653), Mouse anti-pTauT181 (ThermoFisher Scientific, MN1050), Mouse anti-GSK-3βpY216 (BD Bioscience, 612313), Rabbit anti GSK-3β (Cell Signaling, 9315), Mouse anti-ERK1/2 (Cell Signaling, 4696), Rabbit anti-phospho-ERK1/2 MAPK (Cell Signaling, 9101), Rabbit anti-Akt (Ozyme, C67E7), Rabbit anti-phospho-Akt(Ser473) (Ozyme, CST4060), Rabbit anti-p38MAPK (Cell Signaling, 9212), Rabbit anti-phospho-p38MAPK(Thr180/Tyr182) (Cell Signaling, 4511), Rabbit anti-NRXN3 (Abcam, ab192313), and Mouse anti-actin (Sigma, A1979). Secondary antibodies used for the immunoblots were Goat anti-Chicken HRP (ThermoFisher Scientific, A16054), Goat anti-Guinea-pig HRP (Jackson Immunoresearch, 106-035-003), Goat anti-Rabbit HRP (Jackson Immunoresearch, 111-035-003) and Goat anti-Mouse HRP (Jackson Immunoresearch, 115-035-003).

### Alpha-LISA measurements

hNC culture media were collected at 4 weeks of differentiation to dose Aβ using Alpha-LISA assays (Alpha-LISA Amyloid β1–X Kit, AL288C, PerkinElmer) according to the manufacturer’s instructions. 5 μL of cell medium or standard solution was added to an Optiplate-96 microplate (PerkinElmer), incubated with 5 μL of 10X mixture of acceptor beads and biotinylated antibody for 1 h at RT, and then by 40 μL of 1.25X donor beads for 30 min at RT. Luminescence was measured using an EnVision-Alpha Reader (PerkinElmer) at 680 nm excitation and 615 nm emission wavelengths. Aβ concentrations were normalized by the quantity of protein measured using BCA Protein Assay (Thermo Scientific, 23225).

### Microelectrode array recordings and analyses

PNCs were cultured on PDL/Laminin pre-coated MEA 96-well plates (CytoView MEA 96, Axion Biosystems, USA) and extracellular action potentials were recorded at DIV21 in three independent cultures. hNCs were cultured on PLO/Laminin pre-coated MEA 24 or 48-well plates (CytoView MEA 24 or 48, Axion Biosystems, USA) and recorded at DIV28 in three independent cultures using the MaestroPro (Axion Biosystems, Inc, USA). Signals were recorded for 5 min, and we performed multi-unit analysis considering transient potentials detected using 6 standard deviations adaptative threshold crossing methods, and bursts defined as a minimum of 5 spikes in 100 ms. PNCs with mean firing rate <0.2 Hz, hNCs with mean firing rate < 0.1 Hz, and outliers detected with Matlab were excluded from the analysis.

### Morphometric analysis of mouse DG granule cells

10 male control mice (littermates of APP/PS1 mice) aged between 3-4 months old with a body weight of 36.0±1.4 g, were used in this study. In order to silence the plcg2 gene in mice we have injected lentiviral constructs expressing either a shRNA against PLCG2 (CAGTTCCATCTCTTCTATAAA, Vector Builder) or a non-targeting (NT) shRNA (CAACAAGATGAAGAGCACCAA, Vector Builder), combined with GFP expressed under the control of the PGK promoter, in the dorsal dentate gyrus through stereotaxic manipulation. To compare the efficacy of transduction of both LV constructs, we injected in a same mouse LV-plcg2shRNA and LV-scramble in the ipsi- and contra-lateral sides, respectively. After 27 days, mice were transcardially perfused, and the brain was isolated and post-fixed in paraformaldehyde (PFA, 4%). Next, 40-μm-thick sections were prepared with vibratome, immuno-stained and imaged for expression of NeuN and GFP. To analyze the morphology of GFP expressing DG granule cells, we used a confocal microscope (DM5500 TCS SPE, Leica Microsystems) to acquire z-stacks at randomly chosen positions in the supra or infrapyramidal blade of DG. Images were obtained with a 63× NA 1.4 objective and regular photomultipliers. Spine density was measured in four animals with 13 fields at high magnification with >2X zoom in SP1 confocal, at least 2-3 neurites per field. In order to analyze the dendritic and spine morphometry we have used neuronstudio (ver. 0.9.92 64 bit)^65^ and ImageJ^66^ for reconstruction as well as analysis. Subjective investigation of GFP marker co-expressing with the shRNA construct did not show any non-neuronal cell fluorescence up on excitation with 488 nm light source. To check for transduction of microglial cells by the lentiviral construct used for *in-vivo* experiments, we have measured colocalization of IBA1 (with an anti-IBA1 antibody) and GFP immunoreactivity in the contralateral shNT and shPLCG2 injected mice brain.

### *In vivo* viral gene transfer in the DG

Mice (P22–P30) were anesthetized by isoflurane inhalation and injected with buprenorphine to prevent post-surgery pain. 1 μL of either LV-shNT or LV-shPLCG2 solution (5×10^6^ lentiviral particles) was injected using a micropump and syringe (Nanofil WPI) at the rate of 100 nL/min in the DG (Y: 2.2 mm from lambda; X: ± 2.0 mm from sagittal suture; Z: 2.0 mm from the skull) to infect GCs in the dorsal hippocampus. In the acquired images, we have calculated dual positive cells, i.e., IBA1 expressing transfected (GFP+) cells in three animals separately, with contralateral intrahippocampal injection of LV-shNT or LV-shPLCG2 viral constructs using stereotaxic manipulation. For cell counting we have used four random microscopic fields in high magnification. Signals were counted in FIJI using ‘multipoint tool’ in z-projection from image stacks.

### Electrophysiology analysis of mouse DG slices

For electrophysiological recordings, slices were prepared at least 3 weeks post-injection. Mice were anesthetized using ketamine/xylasine mix (100 mg/kg / 10 mg/kg; i.p.) and perfused transcardially with ice cold artificial cerebrospinal fluid (aCSF) for 40–60 s. The brain was quickly removed and placed in ice-cold aCSF containing the following (in mM): 120 NaCl, 26 NaHCO_3_, 2.5 KCl, 1.25 NaH_2_PO_4_, 2 CaCl_2_, 1 MgCl_2_, 16.5 glucose, 2.8 pyruvic acid, 0.5 ascorbic acid, pH 7.4 adjusted by saturating with carbogen (95% O_2_ and 5% CO_2_), and an osmolarity of 305 mOsm. The slices were then transferred to a recording chamber where they were submerged and perfused with an extracellular recording solution containing the following (in mM): 125 NaCl, 2.5 KCl, 1.35 NaH_2_PO_4_, 2 CaCl_2_, 1 MgCl_2_, 16 glucose, 26 NaHCO_3_ continuously oxygenated (95% O_2_ and 5% CO_2_). The recordings were made at RT. Whole-cell patch-clamp recordings (3–4 MΩ electrodes, − 70 mV holding potential) were made from GFP expressing DG granule cells visualized by infrared video-microscopy and epifluorescence. Patch clamp electrodes were pulled out from borosilicate glass (GF 150 F-10). For voltage-clamp recordings, the internal solution contained the following (in mM): 140 CsCH_3_SO_3_, 2 MgCl_2_, 4 NaCl, 5 phospho-creatine, 2 Na_2_ATP, 0.2 EGTA, 10 HEPES, and 0.33 GTP adjusted with CsOH (300 mOsm, pH 7.3). For current-clamp recordings, a K+-based internal solution containing (in mM) was used: 115 K-gluconate, 10 KCl, 0.2 EGTA, 10 HEPES, 15 phospho-creatine, 4 MgATP and 0.3 NaGTP adjusted with KOH (300– 305 mOsm, pH 7.3). The access resistance was <20%. TTX (500 nM) was added to the bath for miniature EPSCs recording. MOhms and the experiment was discarded if it changed by >20%. Recordings were made using an EPC10.0 amplifier (HEKA Elektronik), filtered at 0.5–1 kHz and analysed using IGOR Pro and Neuromatic V2.6 softwares. Evoked AMPA-EPSCs and NMDA-EPSCs were recorded following electrical stimulation of the perforant path (at 0.1 Hz), with an electrode positioned in the molecular layer of the DG, at a membrane potential of −70 mV and +40 mV (in the presence of 10 µM NBQX), respectively. All animal experiments were approved by the local ethical committee of the Institut Pasteur de Lille and according to regulations of the University of Bordeaux/Centre National de la Recherche Scientifique (CNRS) Animal Care and Use Committee.

### Whole exome sequencing data

Written informed consent was obtained from study participants or, for those with substantial cognitive impairment, from a caregiver, legal guardian, or other proxy, and the study protocols for all populations were reviewed and approved by the appropriate local lnstitutional Review Boards^67^. The ADGEN cohort has been collected for a study focusing on the identification of novel Alzheimer’s disease-associated genes and pathways using existing clinical cohorts from Eastern Finland. Research Ethics Committee of Northern Savo, Finland has approved ADGEN study; approval statement 420/13.02.00/2016.

ADGEN cohort comprised 527 AD patients (mean age at onset of AD 72.4 ± 7.9 years, range 43 to 91; 65% women) from Eastern Finland. These AD patients were examined at the Department of Neurology, Kuopio University Hospital and they all fulfilled the NINCDS-ADRDA criteria for probable AD^67^. Whole-exome sequencing (WES) data were generated from ADGEN cohort at the Broad Institute, the Baylor College of Medicine’s Human Genome Sequencing Center, and Washington University’s McDonnell Genome Institute^68^. Multiallelic sites were converted into biallelic variants and then left-aligned using bcftools 1.10. Variants with variant quality score log-odds (VQSLOD) less than −2 or inbreeding coefficient less than −0.3 were excluded. All variants included in final analysis had genotype read depth of at least 10 and genotype quality at least 25^69^. Ensembl Variant Effect Predictor (version v98) (https://www.ensembl.org/info/docs/tools/vep/index.html) was used to predict pathogenicity of PLCG2 variants. To confirm the two nonsense variants in PLCG2 gene identified with WES, DNA was extracted from the blood samples of the carriers using QIAamp DNA Blood Mini Kit (51104, Qiagen). To confirm SNP rs368241065 (C>T, p.R953*), PCR was conducted around the target site by using the following primers: 5′-CACCCACATTGCAACCCTAC-3′, 5′-CTTCGGTCCTCTCATCCCTC-3′. After purification with NucleoSpin® Gel and PCR Clean-up kit (#740609.50, Macherey-Nagel), the PCR product was sequenced at Macrogen Europe (Amsterdam, the Netherlands) with primer 5′-AGAGGGCAAGAGTCCACAG-3′. Novel variant (C>T, p.Q816*) was directed to whole genome sequencing at Novogene Europe. We used the ‘European (Finnish)’ non-neuro subset of the gnomAD v2.1.1 exome dataset as a control cohort (n= 8,707). This cohort contains only samples, which were not collected as part of a neurologic or psychiatric case/control study, or samples that were collected as part of a neurologic or psychiatric case/control study but designated as controls. We identified therein three participants who carried any predicted PLCG2 LoF SNV or indel variants. WES data from Alzheimer Disease European Sequencing (ADES) consortium cohort comprising of 8,732 AD patients and 8,955 control individuals were analyzed by focusing on PLCG2 LoF variants and the same pipeline and QC step were applied as described in^7^. A total of five different PLCG2 LoFs were found in France, The Netherlands and the UK. There were nine carriers of any of the five *PLCG2* LoF variants among 8,732 cases and one carrier among 8,955 controls. All the LoF variants are fully described in supplementary Table S2.

Meta-analysis of the PLCG2 LoF variants in ADGEN and ADES cohorts was performed using the metabin function of the R-package meta (v7.0.0)^70^ using the ‘Mantel-Haenszel’ method and ‘OR’ as the summary measure.

### RNA sequencing in blood

RNA was extracted from the frozen blood of AD patient (age of onset = 60 years) carrying *PLCG2*p.Q816* variant and from four probable AD patients^67^ without PLCG2 LoF variants (mean age of onset = 61±1.4 years; range 60-63) using NucleoSpin® RNA Blood kit (#740200.10, Macherey-Nagel) according to the kit instructions. Library preparation and RNA sequencing was conducted at Novogene Europe (Cambridge, UK) with an Illumina high-throughput sequencing platform. The 150 nt pair-end RNA-seq reads from the 21.7–28.9 million sequenced fragments per sample were quality controlled using FastQC (version 0.11.7) https://www.bioinformatics.babraham.ac.uk/projects/fastqc/). Reads were trimmed with Trimmomatic (version 0.39)^71^ to remove Illumina sequencing adapters and poor quality read ends, using the following essential settings: ILLUMINACLIP:2:30:10:2:true, SLIDINGWINDOW:4:10, LEADING:3, TRAILING:3, MINLEN:50. The trimmed reads were aligned to the Gencode human transcriptome version 38 (for genome version hg38) using STAR (version 2.7.9a)^72^ with essential non-default settings: - seedSearchStartLmax 12, - alignSJoverhangMin 15, - outFilterMultimapNmax 100, - outFilterMismatchNmax 33, - outFilterMatchNminOverLread 0, - outFilterScoreMinOverLread 0.3, and - outFilterType BySJout. The unstranded, uniquely mapping, gene-wise counts for primary alignments were collected in R (version 4.3.2) using Rsubread::featureCounts (version 2.16.1)^73^, totaling 14.2 to 18.0 million per sample. Read counts for PLCG2 gene were normalized for differences in sequencing depth using size factors calculated by estimateSizeFactors (from DESeq2 version 1.38.3)^74^.

### AD hallmarks in human brain and CSF samples

Human post-mortem brain samples were originally obtained from Kuopio University Hospital and investigated within Memory polyclinic research projects. Following autopsy, brains were fixed in buffered 10 % formalin for at least one week. Tissue blocks were prepared, embedded in paraffin, and used for evaluation of AD-related pathology and classified according to Braak staging as described before^75^. Detailed information of the cohort and handling of the samples have been provided in several previous publications^76–78^. Individual carrying the p.R953* variant in *PLCG2* was analyzed together with five matched controls. All analyzed subjects were females (mean age 84.3±4.03), APOE ε3/4, diagnosed with clinical dementia, and defined as Braak stage VI in the neuropathological examination.

### Immunohistochemistry

Paraffin embedded inferior temporal lobe samples were serially sectioned in 7 µm thickness. The sections were deparaffinized in xylene and rehydrated in graded alcohol series. For IBA1/Aβ co-staining, sections were processed and stained with rabbit anti-IBA1 (1:1000, #019– 19,741, FUJIFILM Wako Chemicals Europe GmbH) and mouse anti-Aβ (1:500, anti-β-amyloid 17–24, clone 4G8, #800712, 1:500, Biolegend) following the procedure explained elsewhere^79^.

For fluorescent staining, antigen retrieval was performed in 10 mM Tris – 1 mM EDTA buffer, pH 9.0 in pressure cooker for 10 min for all sections except those used for p-Tau staining. Additional antigen retrieval step in 80 % formic acid (20 min, RT) was carried out for sections used for CD68/Aβ/IBA1 staining. Autofluorescence was quenched using TrueBlack lipofuscin autofluorescence quencher (#23007, Biotium). Target proteins were visualized by following primary antibodies: goat anti-Apolipoprotein E (1:1000, #178479, Merck Millipore), goat-anti IBA1 (1:1000, #ab5076, Abcam), mouse anti-Aβ (1:500, anti-β-amyloid 17–24, clone 4G8, #800712, Biolegend), rabbit anti-GFAP (1:1000, #Z0334, Dako), rabbit anti-P2RY12 (1:1000, #HPA014518, Sigma Aldrich), and mouse anti-phospho-tau detecting phosphorylated Ser202, Thr205, and Ser208 residues (1:1000, B6, produced in house^79–81^) coupled with appropriate species-specific secondary antibodies: donkey anti-goat IgG Alexa Fluor 568 (1:500, #A-11057, Invitrogen), donkey anti-goat IgG Alexa Fluor 633 (1:500, #A21082, Invitrogen), goat anti-mouse IgG Alexa Fluor 488 (1:2000, #A-11001, Invitrogen), donkey anti-mouse IgG Alexa Fluor 568 (1:500, #A10037, Invitrogen), goat anti-rabbit IgG Alexa Fluor 680 (1:500, #A-21109, Invitrogen). CD68 was visualized by rabbit anti-CD68 (1:5000, #ab213363, Abcam) followed by horse anti-rabbit biotinylated IgG (1:200, BA-1100, Vector Laboratories) and Alexa Fluor 488 Tyramide SuperBoost streptavidin kit (B40932, Invitrogen). To visualize β-sheet structures within Aβ plaques, sections were incubated 10 min in 100 µM X-34 (1,4-Bis(3-carboxy-4-hydroxyphenylethenyl)benzene, Sigma-Aldrich) in 40% ethanol, and differentiated in 80% ethanol with 0.2g% NaOH for 2 min. Nuclei were visualized with DAPI (1 µg/ml) and the sections were embedded with Vectashield Hardset or Vectashield Vibrance antifade mounting medium (H-1400 and H-1700, Vector Laboratories).

### Imaging and analysis of human brain tissue

IBA1/Aβ co-stained sections were imaged with Hamamatsu NanoZoomer-XR Digital slide scanner with 20× (NA 0.75) objective (Hamamatsu Photonics K.K.). NDP.view2 software (Hamamatsu Photonics K.K.) was used for counting the number and the size of β-amyloid plaques and the number of β-amyloid plaque-associated microglia. For analysis, seven 250 µm2 blocs were randomly chosen from the grey matter area in each brain section. All plaques within the blocks were included in the analysis. Size of the β-amyloid plaques was obtained by manually outlining the plaques. Plaque-associated microglia (within or touching the plaque) were manually counted. Only microglia with clearly visible soma were included in the count. Counting and analysis was performed by investigators blinded to sample identity.

P2RY12, GFAP, and B6 immunofluorescence stainings were imaged with THUNDER Imager 3D Tissue imaging system equipped with 20× objective and using the built-in computational clearing method to remove out-of-focus signal (Leica Microsystems). Seven to ten randomly selected fields of view in cortical grey matter were imaged in each sample. Images were further processed with Fiji^66^ by applying background subtraction (Sliding paraboloid, Rolling ball radius 50 pixels) and Gaussian blur filtering before masking the images using the automatic threshold (Li algorithm for GFAP, Triangle for P2RY12, Moments for B6, and Li for DAPI) and measuring the immunopositive area. APOE, X-34, CD68, IBA1, and 4G8 immunofluorescence stainings were imaged with Axio Observer inverted microscope equipped with a LSM800 confocal module with Plan-Apochromat 20× or 40× (NA 0.8) objective (Carl Zeiss Microimaging GmbH). A z-stack was captured from six to eight randomly selected fields of view. A sum projection of the stack was created in Fiji, after which the background was subtracted (Sliding paraboloid, Rolling ball radius 50 pixels) and Gaussian blur filtering applied. The images were then masked using the automatic threshold algorithm (Triangle for APOE, IBA1, and 4G8, Moments for X-34 and CD68), and immunopositive area was measured. All imaging and analyses were performed by investigators blinded to sample identity.

### Processing and analysis of frozen brain tissue samples

Frozen brain tissue from the same individuals were homogenized, and proteins were extracted as described previously^76^. The protein concentration of the lysates was measured with the Pierce BCA Protein Assay Kit (#23227, Thermo Fisher Scientific). 18 µg of total protein per sample were denatured at 55°C for 10 min with NuPAGE LDS Sample Buffer (#NP0007, Invitrogen). Proteins were separated on NuPAGE 4–12% BisTris Midi Protein Gels (#WG1202BOX, Invitrogen) and transferred onto PVDF membranes (#IB24001, Thermo Fisher Scientific) with an iBlot 2 device (IB21001, Thermo Fisher Scientific). Blot was probed overnight at +4°C with PLCγ2 (1:200, clone B-10, #sc-5283, Santa Cruz) and β-actin (1:1000, #ab8226, Abcam) antibodies. After incubation with mouse IgG horseradish peroxidase linked secondary antibody (1:5000, #GENA931, Merck) for 1 h, proteins were detected with ECL Select Western Blotting Detection Reagent (#12644055, Merck) and a ChemiDoc Imaging System (Bio-Rad Laboratories). Image Lab Software 6.0.1 (Bio-Rad Laboratories) was used to quantify protein levels. The levels of PLCγ2 were normalized to that of β-actin within each sample. Aβ_42_ levels were measured from soluble fraction using HRP-conjugated antibody-based Human/Rat β-Amyloid 42 (High Sensitive, #29062601, Wako) ELISA Kit as has been described before^78^. Aβ_42_ concentrations were normalized to total protein concentration of the same sample.

#### CSF analyses

Aβ_42_, total-tau (tau), and phosphorylated-tau (p-tau) levels in the CSF were measured using Innotest β-amyloid 1-42, Innotest hTau Ag, and Innotest Phospho-tau 181P ELISA kits (Innogenetics, Ghent, Belgium) according to the protocol described previously^77,78^.

### Analysis of single-nucleus RNA-sequencing datasets of human brains

We obtained two publicly available datasets of single-nucleus RNA-seq on human brain tissue, published by Li et al. 2025^22^ (GSE237718 from GEO) and Leng et al. 2021^23^ (syn21788402 from Synapse). The former was performed on temporal cortex tissue, and the latter was performed on entorhinal cortex and superior frontal gyrus. Analysis was performed using the R package Seurat^82^. For Li et al. 2025^22^, each individual’s data was read, keeping cells with at least 200 detected features and fraction of mitochondrial genes less than 10%, and keeping features detected in at least 10 cells. Data were SCTransformed while regressing on percent mitochondrial genes^83^. FindNeighbors was run using 30 principal component analysis (PCA) dimensions, and FindClusters with resolution 0.35. Clusters were classified into the following cell types using a set of rules based on marker genes: microglia (PTPRC), astrocytes (SLC1A2), oligodendrocytes (PLP1), oligodendrocyte precursor cells (VCAN), inhibitory neurons (GAD1 minus VCAN-positive clusters), excitatory neurons (CAMK2A minus GAD1-positive clusters), and vascular/endothelial cells (CLDN5). For Leng et al. 2021^23^, cells were already annotated with cell types.

### Single-nucleus RNA-sequencing of hNCs cultures

#### Nuclei isolation

The isolation of hNCs nuclei was conducted on ice using pre-cold buffers, followed by centrifugation at 500 rcf and 4°C for 15 min. First, cells were resuspended in lysis buffer: 10 mM Tris-HCl, 10 mM NaCl, 3 mM MgCl_2_, 0.1% NP40 and 0.04 U/μl RNase OUT (Invitrogen, 10777019). The nuclei pellet was subsequently washed three times with nuclei wash buffer (1× DPBS, 1% BSA, and 0.04 U/μl RNase OUT). The cells were then filtered through a 40 µm Flowmi strainer (Merck, BAH136800040-50EA) and fixed using the Single Cell Fixed RNA Sample Preparation Kit (10X genomics, PN-1000414). The nuclei were preserved in 10X nuclei preservation buffer and stored at −80°C until four independent differentiations were completed.

#### Library preparation and sequencing

The day of the hybridization protocol, samples from all four conditions were thawed and processed in parallel. Each experimental condition (shNT, shPLCG2, WT and R953*) was hybridized using one of the four probes from the Chromium Next GEM Single Cell Fixed RNA Human Transcriptome Probe Kit (PN-1000420; 10X Genomics). Samples were then pooled to create a single mix containing all four conditions and to target 40,000 encapsulated nuclei per lane (n=4 lanes) according to the manufacturer’s protocol. Pooled nuclei were then encapsulated by using Single Cell TL v1 Gel Bead (PN-2000538; 10X Genomics) and Next GEM Chip Q (PN-2000518; 10X Genomics) on the Chromium iX. SnRNA-seq libraries were prepared by using the PTC-200 Thermal Cycler (MJ Research) and by following the Fixed RNA Profiling Reagent Kit’s user guide (CG000527 Rev F; 10X Genomics). Library quality was assessed with the Bioanalyzer 2100 (Agilent) and their cDNA quantity was assayed using the Quant-IT^TM^ 1X dsDNA HS Assay Kit (Ref Q33232; Invitrogen). All four libraries were combined in equimolar quantities for a single sequencing pool. This pool was sequenced (BGI, Hong Kong) according to the manufacturer’s instructions (28:10:10:90, R1:i7:i5:R2) targeting a depth of ∼60k reads/nuclei.

#### snRNA-seq data preprocessing and annotation

Sequencing reads for the four multiplexed libraries have been processed using Cellranger Multi (10X Genomics) in Multiplex Flex mode using associated human genome reference refdata-gex-GRCh38-2024-A and Human Transcriptome probes set v1.1.0_GRCh38-2024-A, calling in average 15k cells per sample, median of 2500 genes and 4000 UMis detected per cell. We then used Seurat (V5.2.1)^82^ to process the generated gene expression matrices. To remove low quality and aggregated nuclei, we removed all nuclei with more than 8% mitochondrial count or number of total count exceeding 100k. In addition, we removed nuclei with both low gene detection (less than 1500) and having high mitochondrial RNA percentage (greater than 2%), likely reflecting high mitochondrial contamination over degraded nuclei (supplementary Fig S8a,b). We observed at this stage that one replicate of shPLCG2 (shPLCG2_4) had a strong mitochondrial percentage (median = 3% while all other samples had <1%) and low RNA and gene counts; we thus excluded this replicate as well as the associated control (shNT_4) from the analysis. Then, the gene expression matrices were normalized using Seurat::SCTransform() with default parameters but regressing out for mitochondrial percentage. PCA was performed on the residuals of the 3000 most variable genes, and harmony::RunHarmony() (1.2.0)^84^ was used to reduce influence of batch effect on PCs by correcting for sample specific variability. The 20 first harmony corrected component was used to generate a Shared Nearest Neighbor graph using Seurat::FindNeighbors() and cluster nuclei on this graph with Louvain algorithm using Seurat::FindClusters() with resolution of 0.5. UMAP with Seurat::RunUMAP() was then performed to generate a two-dimension space non linearly representing the gene expression variability between nuclei. A total of 23 clusters were identified (supplementary Fig S8c) with mix of samples within each cluster (supplementary Fig. S8d, g). These clusters were annotated based on canonical neuronal markers expression in each cluster (supplementary Fig. S8f). From these 23 clusters, we identified: Astrocytes, subdivided into 4 subpopulations (steady state: clusters 1, 6, 7 and 18; proliferative: clusters 4 and 8; immature: cluster 10; with VLMC potential: clusters 13 and 22); Excitatory Neurons (clusters 12, 19 and 20); Inhibitory Neurons (clusters 5, 14 and 17); NPCs (cluster 9); Vascular leptomeningeal cells (VLMC, cluster 11); and Fibroblasts (cluster 21). Cluster 0 contained a mix of excitatory and inhibitory neurons and was therefore subdivided into two clusters based on the expression of glutamate transporter SLC17A6 and the GABAergic enzymes GAD1/2. Cluster 16, expressing markers of more than one cell type and with greater RNA content than others, was deemed as doublet cells and removed. Clusters 2, 3 and 15 having low RNA content, high mitochondrial RNA percentage, low cell type identity and/or expressing stress markers (Chaperone/ heat shock protein gene family, stress granules TIA1) were filtered out as well (supplementary Fig. S8e). The resulting filtered matrix and clusters alongside with the raw feature barcode matrices were used to estimate ambient RNA contamination in each sample using SoupX (1.6.2)^85^. Probes based snRNA-seq assay having higher gene detection but also higher ambient RNA level compared to standard polyA based mRNA capture technic, we increased stringency in determining contamination threshold using for each sample SoupX::autoEstCont() with tfidfMin=.95, soupQuantile =0.6, priorRho = 0.1, and contaminationRange = c(0.05,0.8) parameter to automatically estimate ambient RNA level. The estimated ambient level was 14% on average (min: 5.1%, max: 34%) across samples. We then used SoupX::adjustCounts() to correct the gene expression matrix accordingly, and we further ensured no impact on downstream differential analysis using the ambient RNA level as covariate.

#### Neuronal maturation score calculation

We calculated a neuronal maturation score using a published neuron-specific maturation gene set (MS-NS, 25 genes)^86^ and performed cell level enrichment analysis using UCell^87^, AddModuleScore_UCell function with parameter maxRank=1500. We then computed the average score per donor and cell type.

#### Differential expression and pathway enrichment analysis

Cell type-specific differential expression analyses comparing R953* *vs* WT and shPLCG2 *vs* shNT were performed using DESeq2(v1.40.2)^74^ and the SoupX corrected-count matrix. For each cell type, the genes were filtered to keep only those for which more than 100 nuclei or 10% of the nuclei have at least 5 counts. The counts were then aggregated per sample, and the resulting pseudobulk count matrix was further filtered out by keeping only genes with more than 50 CPM in at least 20% of the samples. Negative binomial model-based regression using DESeq2 was then performed to estimate effect of the treatment or genotype, correcting for average mitochondrial RNA percentage, ambient RNA contamination rate, and library batch. Before running DESeq2::DESeq() model fitting, the size factors (from DESeq2::estimateSizeFactors()) were estimated using the unfiltered pseudobulk matrix. Were considered as DEGs, genes with FDR <0.05 and |log2FC|>0.5. Functional enrichment analysis was performed on these DEGs using hypergeometric test (base::phyper()) and MSigDB Gene Ontology and Canonical Pathways (c2.cp.v2023 and C5.go.v2023).

#### Regulon analysis

We performed the TF regulon identification using pyscenic 0.12.1.^88^(with grnboost2) on 20,000 randomly sampled nuclei transcriptome previously normalized and log-transformed using sc.pp.normalize total and sc.pp.log1p.^89^ We used ‘hg38_10kbp_up_10kbp_down_full_tx_v10’ genes-motifs ranking and ‘motifs-v10nr’ TF motifs from cis target database https://resources.aertslab.org/cistarget/databases/homo_sapiens/hg38/refseq_r80/mc_v10_clust/gene_based/. We then computed the cell level activity score of the identified TF regulons on the full dataset using pyscenic.aucell. The resulting score for each regulon was then averaged per sample and cell type, and Wilcoxon ranks sum test was performed to compare the conditions on selected regulons.

#### Cell-Cell Communication

Cell-Cell communication with ligand receptor (L/R) interaction inference was performed using CellChat (2.2.0)^34^. For each condition, replicates were first grouped into one CellChat object and communication probabilities between cell types were determined using CellChatDB.human reference excluding non-protein signaling. CellChat::identifyOverExpressedGenes, CellChat::identifyOverExpressedInteractions and CellChat::computeCommunProb were used with default parameters to measure probability of communication (interaction strength and statistical significance) based on ligand, receptor and cofactors overexpression. To filter L/R not conserved across biological replicates, CellChat::filterCommunication was used with min.samples = 3 and min.cells = 10. These measurements were aggregated at signaling pathways level using CellChat::computeCommunProbPathway. Overall incoming and outgoing signaling L/R and signaling pathway interaction strength for each cell type were then computed using CellChat::netAnalysis_computeCentrality. Difference of strength between treated and respective control condition was determined to obtain the differential incoming/outgoing interaction, where difference of strength > 0.25 was considered significant. To further confirm robustness of the differences observed across condition for the significant signaling pathways and L/R interaction, cell-cell communication inference was performed again for each sample separately, and the interaction strength per cell type was collected for each sample and compared between conditions using Wilcoxon rank sum test.

#### Comparison with dementia DEGs

SEA-AD DLPFC and MTG snRNA-seq datasets were downloaded from https://registry.opendata.aws/allen-sea-ad-atlas/. Donors annotated as ‘Reference’ (lacking cognitive status), and with non-Caucasian ethnicity have been excluded from analysis, as well as donors with abnormal cell type distribution. Abnormal cell type distribution was measured by calculating the cell type proportion deviation from the interquartile range (IQR) and by computing the PCA of these deviation matrices. Donors with first principal component coordinates at 3 times away from the IQR were excluded from analysis. A total of 12 donors were excluded for both DLPFC and MTG regions. For each cell type, donors with more than 50 nuclei were considered. A donor level count matrix was then generated by aggregating nuclei count by donor (total number of donors kept for analysis = 70). The differential expression was performed on comparing the individuals with dementia compared to individuals without dementia using DESeq2, adjusting for sex, post mortem interval (PMI), atherosclerosis stage, library input (ng), number of PCR cycles, number of cells, median number of molecules detected per cell, average mitochondrial RNA percentage.

---

**Extended Figure 1:**
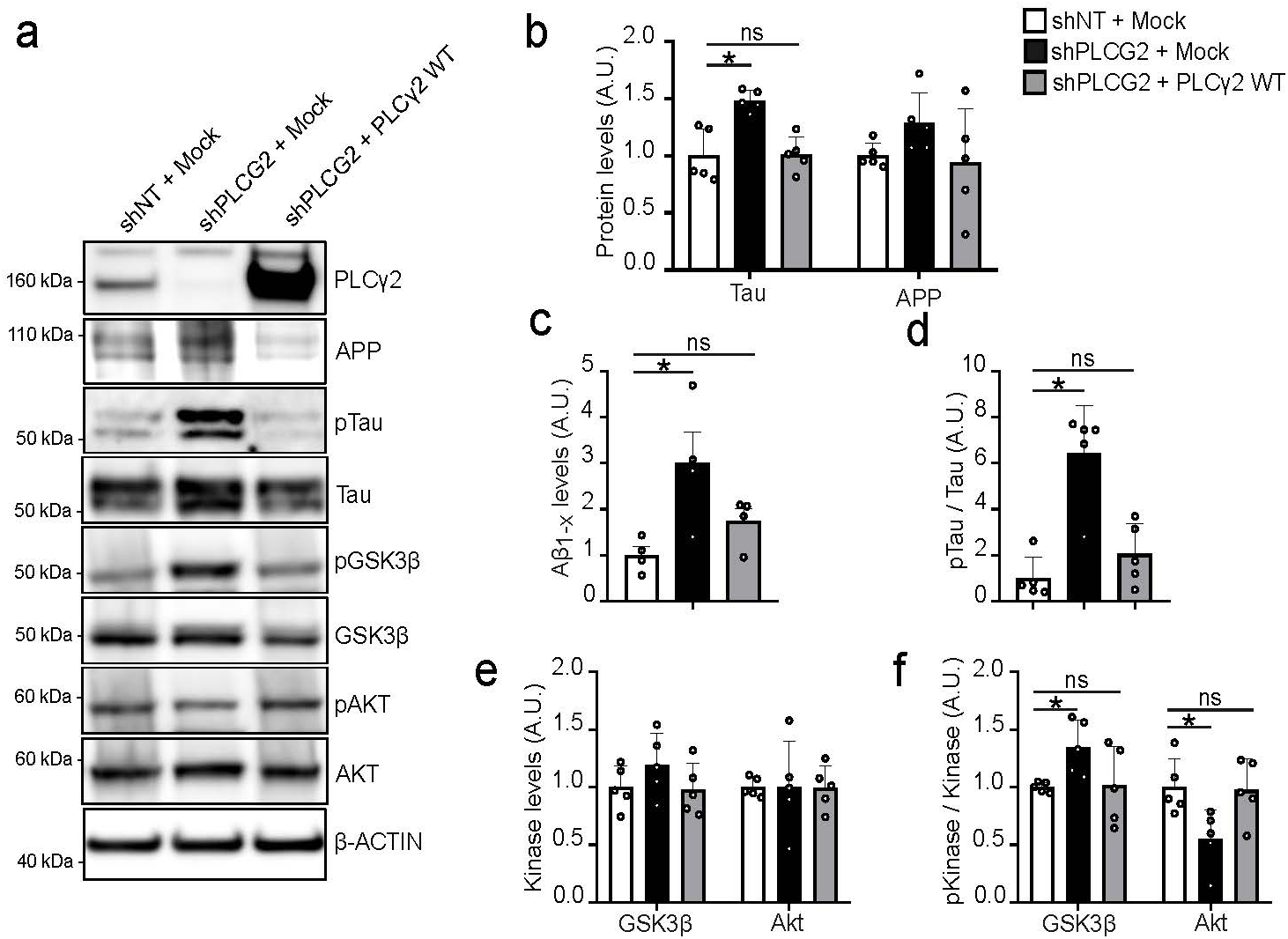
PLCG2 re-expression in shPLCG2-hNCs reverse AD hallmark changes. (a) Representative Western blots of PLCγ2, APP, total and T231-phospho-Tau, total and phospho-GSK3β, and total and phospho-AKT protein levels in shNT-hNCs, shPLCG2-hNCs, and shPLCG2-hNCs overexpressing WT PLCG2 (shPLCG2-hNCs + PLCG2). (b) Quantification of APP and total Tau protein levels in shNT-hNCs, shPLCG2-hNCs, and shPLCG2-hNCs + PLCG2. Data given as mean±SD, n=4. (c) Aβ peptide levels in the culture medium, measured *via* Alpha-LISA and normalized by total protein levels, in shNT-hNCs, shPLCG2-hNCs, and shPLCG2-hNCs + PLCG2. Data given as mean±SEM, n=4. (d) Quantification of relative Tau phosphorylation at T231 in shNT-hNCs, shPLCG2-hNCs, and shPLCG2-hNCs + PLCG2. Data given as mean±SD, n=4. (e-f) Changes in total protein levels (e) and relative phosphorylation (f) of GSK3β and AKT in shNT-hNCs, shPLCG2-hNCs, and shPLCG2-hNCs + PLCγ2. Data given as mean±SD, n=4. All statistical analyses were performed using Kruskal-Wallis test followed by Dunn’s multiple comparison test. *p<0.05.

## REFERENCES

1. van Dyck, C. et al. Lecanemab in Early Alzheimer’s Disease. N. Engl. J. Med. 388, 142–143 (2023).

2. Herrup, K. The case for rejecting the amyloid cascade hypothesis. Nat. Neurosci. 18, 794–799 (2015).

3. Selkoe, D. J. & Hardy, J. The amyloid hypothesis of Alzheimer’s disease at 25 years. EMBO Mol. Med. 8, 595–608 (2016).

4. Frisoni, G. B. et al. The probabilistic model of Alzheimer disease: the amyloid hypothesis revised. Nat. Rev. Neurosci. 23, 53–66 (2022).

5. Gatz, M. et al. Role of Genes and Environments for Explaining Alzheimer Disease. Arch. Gen. Psychiatry 63, 168 (2006).

6. Bellenguez, C. et al. New insights into the genetic etiology of Alzheimer’s disease and related dementias. Nat. Genet. 54, 412–436 (2022).

7. Holstege, H. et al. Exome sequencing identifies rare damaging variants in ATP8B4 and ABCA1 as risk factors for Alzheimer’s disease. Nat. Genet. 54, 1786–1794 (2022).

8. Lambert, J. C., Ramirez, A., Grenier-Boley, B. & Bellenguez, C. Step by step: towards a better understanding of the genetic architecture of Alzheimer’s disease. Mol. Psychiatry 28, 2716– 2727 (2023).

9. Schürmann, B. et al. A novel role for the late-onset Alzheimer’s disease (LOAD)-associated protein Bin1 in regulating postsynaptic trafficking and glutamatergic signaling. Mol. Psychiatry 25, 2000–2016 (2020).

10. Eysert, F. et al. Alzheimer’s genetic risk factor FERMT2 (Kindlin-2) controls axonal growth and synaptic plasticity in an APP-dependent manner. Mol. Psychiatry 26, 5592–5607 (2021).

11. Giralt, A. et al. PTK2B/Pyk2 overexpression improves a mouse model of Alzheimer’s disease. Exp. Neurol. 307, 62–73 (2018).

12. Ojelade, S. A. et al. cindr, the Drosophila Homolog of the CD2AP Alzheimer’s Disease Risk Gene, Is Required for Synaptic Transmission and Proteostasis. Cell Rep. 28, 1799–1813.e5 (2019).

13. Masliah, E. Mechanisms of synaptic dysfunction in Alzheimer’s disease. Histol. Histopathol. 10, 509–519 (1995).

14. Barthet, G. & Mulle, C. Presynaptic failure in Alzheimer’s disease. Prog. Neurobiol. 194, 101801 (2020).

15. Ribarič, S. Detecting Early Cognitive Decline in Alzheimer’s Disease with Brain Synaptic Structural and Functional Evaluation. Biomedicines 11, 355 (2023).

16. Coulon, A., et al. High-Content Screening of Synaptic Density Modulators in Primary Neuronal Cultures. Curr. Protoc. 3, e904 (2023).

17. Kilinc, D. et al. Pyk2 overexpression in postsynaptic neurons blocks amyloid β1-42-induced synaptotoxicity in microfluidic co-cultures. Brain Commun. 2, fcaa139 (2020).

18. Takalo, M. et al. The Alzheimer’s disease-associated protective Plcγ2-P522R variant promotes immune functions. Mol. Neurodegener. 15, 1–14 (2020).

19. Claes, C. et al. The P522R protective variant of PLCG2 promotes the expression of antigen presentation genes by human microglia in an Alzheimer’s disease mouse model. Alzheimer’s Dement. 18, 1765–1778 (2022).

20. Tsai, A. P. et al. PLCG2 is associated with the inflammatory response and is induced by amyloid plaques in Alzheimer’s disease. Genome Med. 14, 17 (2022).

21. Tsai, A. P. et al. Genetic variants of phospholipase C-γ2 alter the phenotype and function of microglia and confer differential risk for Alzheimer’s disease. Immunity 56, 2121–2136. (2023).

22. Li, Z. et al. APOE genotype determines cell-type-specific pathological landscape of Alzheimer’s disease. Neuron 113, 1380–1397. (2025).

23. Leng, K. et al. Molecular characterization of selectively vulnerable neurons in Alzheimer’s disease. Nat. Neurosci. 24, 276–287 (2021).

24. Magno, L. et al. Alzheimer’s disease phospholipase C-gamma-2 (PLCG2) protective variant is a functional hypermorph. Alzheimers. Res. Ther. 11, 16 (2019).

25. Jiang, N. et al. Impaired plasticity of intrinsic excitability in the dentate gyrus alters spike transfer in a mouse model of Alzheimer’s disease. Neurobiol. Dis. 154, 105345 (2021).

26. Viana Da Silva, S., et al. Hippocampal Mossy Fibers Synapses in CA3 Pyramidal Cells Are Altered at an Early Stage in a Mouse Model of Alzheimer’s Disease. J. Neurosci. 39, 4193–4205 (2019).

27. Saha, O. et al. The Alzheimer’s disease risk gene BIN1 regulates activity-dependent gene expression in human-induced glutamatergic neurons. Mol. Psychiatry 29, 2634–2646 (2024).

28. Lejeune, F. Nonsense-Mediated mRNA Decay, a Finely Regulated Mechanism. Biomedicines 10, 141 (2022).

29. Rekha, A. et al. GSK-3β dysregulation in aging: Implications for tau pathology and Alzheimer’s disease progression. Mol. Cell. Neurosci. 133, 104005 (2025).

30. De Simone, A., Tumiatti, V., Andrisano, V. & Milelli, A. Glycogen Synthase Kinase 3β: A New Gold Rush in Anti-Alzheimer’s Disease Multitarget Drug Discovery? J. Med. Chem. 64, 26–41 (2021).

31. Lauretti, E., Dincer, O. & Praticò, D. Glycogen synthase kinase-3 signaling in Alzheimer’s disease. Biochim. Biophys. Acta Mol. Cell Res. 1867, 118664 (2020).

32. Huo, X. et al. GSK3 protein positively regulates type I insulin-like growth factor receptor through forkhead transcription factors FOXO1/3/4. J. Biol. Chem. 289, 24759–24770 (2014).

33. Inestrosa, N. C. & Varela-Nallar, L. Wnt signaling in the nervous system and in Alzheimer’s disease. J. Mol. Cell Biol. 6, 64–74 (2014).

34. Jin, S., Plikus, M. V. & Nie, Q. CellChat for systematic analysis of cell–cell communication from single-cell transcriptomics. Nat. Protoc. 20, 180–219 (2025).

35. Martinez-Mir, A. et al. Genetic study of neurexin and neuroligin genes in Alzheimer’s disease. J. Alzheimers. Dis. 35, 403–412 (2013).

36. Rahman, M. M., Westermark, G. T., Zetterberg, H., Härd, T. & Sandgren, M. Protofibrillar and Fibrillar Amyloid-β Binding Proteins in Cerebrospinal Fluid. J. Alzheimer’s Dis. 66, 1053–1064 (2018).

37. Lee, A. K., Khaled, H., Chofflet, N. & Takahashi, H. Synaptic Organizers in Alzheimer’s Disease: A Classification Based on Amyloid-β Sensitivity. Front. Cell. Neurosci. 14, 567721 (2020).

38. Hishimoto, A. et al. Neurexin 3 transmembrane and soluble isoform expression and splicing haplotype are associated with neuron inflammasome and Alzheimer’s disease. Alzheimers. Res. Ther. 11, 28 (2019).

39. Bot, N., Schweizer, C., Halima, S. Ben & Fraering, P. C. Processing of the synaptic cell adhesion molecule neurexin-3beta by Alzheimer disease alpha- and gamma-secretases. J. Biol. Chem. 286, 2762–2773 (2011).

40. Gabitto, M. I. et al. Integrated multimodal cell atlas of Alzheimer’s disease. Nat. Neurosci. 27, 2366–2383 (2024).

41. Reddy, P. H. Amyloid beta-induced glycogen synthase kinase 3β phosphorylated VDAC1 in Alzheimer’s disease: implications for synaptic dysfunction and neuronal damage. Biochim. Biophys. Acta 1832, 1913–1921 (2013).

42. Hernandez, F., Lucas, J. J. & Avila, J. GSK3 and tau: two convergence points in Alzheimer’s disease. J. Alzheimers. Dis. 33 **Suppl 1**, S141–4 (2013).

43. Ly, P. T. T. et al. Inhibition of GSK3β-mediated BACE1 expression reduces Alzheimer-associated phenotypes. J. Clin. Invest. 123, 224–235 (2013).

44. Ochalek, A. et al. Neurons derived from sporadic Alzheimer’s disease iPSCs reveal elevated TAU hyperphosphorylation, increased amyloid levels, and GSK3B activation. Alzheimer’s Res. Ther. 9, 90 (2017).

45. Sita, L. V., Diniz, G. B., Horta, J. A. C., Casatti, C. A. & Bittencourt, J. C. Nomenclature and comparative morphology of the teneurin/TCAP/ADGRL protein families. Front. Neurosci. 13, 425 (2019).

46. Bourgeron, T. A synaptic trek to autism. Curr. Opin. Neurobiol. 19, 231–234 (2009).

47. Vitobello, A. et al. ADGRL1 haploinsufficiency causes a variable spectrum of neurodevelopmental disorders in humans and alters synaptic activity and behavior in a mouse model. Am. J. Hum. Genet. 109, 1436–1457 (2022).

48. Cvetkovska, V. et al. Neurexin-β Mediates the Synaptogenic Activity of Amyloid Precursor Protein. J. Neurosci. 42, 8936–8947 (2022).

49. Kleineidam, L. et al. PLCG2 protective variant p.P522R modulates tau pathology and disease progression in patients with mild cognitive impairment. Acta Neuropathol. 139, 1025–1044 (2020).

50. Mangiolaa, S. et al. Sccomp: Robust differential composition and variability analysis for single-cell data. Proc. Natl. Acad. Sci. U. S. A. 120, e2203282120 (2023).

## REFERENCES

51. Kaech, S. & Banker, G. Culturing hippocampal neurons. Nat. Protoc. 1, 2406–2415 (2006).

52. Pantazis, C. B. et al. A reference human induced pluripotent stem cell line for large-scale collaborative studies. Cell Stem Cell 29, 1685–1702. (2022).

53. Duncan, G. T., Balamurugan, K., Budowle, B., Smerick, J. & Tracey, M. L. Microvariation at the human D1S80 locus. Int. J. Legal Med. 110, 150–154 (1997).

54. Weisheit, I. et al. Simple and reliable detection of CRISPR-induced on-target effects by qgPCR and SNP genotyping. Nat. Protoc. 16, 1714–1739 (2021).

55. Garcia, M. et al. Sarek: A portable workflow for whole-genome sequencing analysis of germline and somatic variants. F1000Research 9, 63 (2020).

56. Rausch, T. et al. DELLY: structural variant discovery by integrated paired-end and split-read analysis. Bioinformatics 28, i333–i339 (2012).

57. Chen, X. et al. Manta: rapid detection of structural variants and indels for germline and cancer sequencing applications. Bioinformatics 32, 1220–1222 (2016).

58. Pedersen, B. S. & Quinlan, A. R. Duphold: scalable, depth-based annotation and curation of high-confidence structural variant calls. Gigascience 8, giz040 (2019).

59. Jeffares, D. C. et al. Transient structural variations have strong effects on quantitative traits and reproductive isolation in fission yeast. Nat. Commun. 8, 14061 (2017).

60. Concordet, J. P. & Haeussler, M. CRISPOR: Intuitive guide selection for CRISPR/Cas9 genome editing experiments and screens. Nucleic Acids Res. 46, W242–W245 (2018).

61. Chambers, S. M. et al. Highly efficient neural conversion of human ES and iPS cells by dual inhibition of SMAD signaling. Nat. Biotechnol. 27, 275–280 (2009).

62. Bardy, C. et al. Neuronal medium that supports basic synaptic functions and activity of human neurons in vitro. Proc. Natl. Acad. Sci. U. S. A. 112, E2725–E2734 (2015).

63. Kunkle, B. W. et al. Genetic meta-analysis of diagnosed Alzheimer’s disease identifies new risk loci and implicates Aβ, tau, immunity and lipid processing. Nat. Genet. 51, 414–430 (2019).

64. Blasiak, A., Lee, G. U. & Kilinc, D. Neuron Subpopulations with Different Elongation Rates and DCC Dynamics Exhibit Distinct Responses to Isolated Netrin-1 Treatment. ACS Chem. Neurosci. 6, 1578–1590 (2015).

65. Rodriguez, A., Ehlenberger, D. B., Hof, P. R. & Wearne, S. L. Rayburst sampling, an algorithm for automated three-dimensional shape analysis from laser scanning microscopy images. Nat. Protoc. 1, 2152–61 (2006).

66. Schindelin, J. et al. Fiji: an open-source platform for biological-image analysis. Nat. Methods 9, 676–82 (2012).

67. McKhann, G. et al. Clinical diagnosis of Alzheimer’s disease: report of the NINCDS-ADRDA Work Group under the auspices of Department of Health and Human Services Task Force on Alzheimer’s Disease. Neurology 34, 939–944 (1984).

68. Bis, J. C. et al. Whole exome sequencing study identifies novel rare and common Alzheimer’s-Associated variants involved in immune response and transcriptional regulation. Mol. Psychiatry 25, 1859–1875 (2020).

69. Korpioja, A. et al. Novel Rare SORL1 Variants in Early-Onset Dementia. J. Alzheimers. Dis. 82, 761–770 (2021).

70. Balduzzi, S., Rücker, G. & Schwarzer, G. How to perform a meta-analysis with R: a practical tutorial. Evid. Based. Ment. Health 22, 153–160 (2019).

71. Bolger, A. M., Lohse, M. & Usadel, B. Trimmomatic: a flexible trimmer for Illumina sequence data. Bioinformatics 30, 2114–2120 (2014).

72. Dobin, A. et al. STAR: ultrafast universal RNA-seq aligner. Bioinformatics 29, 15–21 (2013).

73. Liao, Y., Smyth, G. K. & Shi, W. The R package Rsubread is easier, faster, cheaper and better for alignment and quantification of RNA sequencing reads. Nucleic Acids Res. 47, e47 (2019).

74. Love, M. I., Huber, W. & Anders, S. Moderated estimation of fold change and dispersion for RNA-seq data with DESeq2. Genome Biol. 15, 550 (2014).

75. Braak, H., Alafuzoff, I., Arzberger, T., Kretzschmar, H. & Tredici, K. Staging of Alzheimer disease-associated neurofibrillary pathology using paraffin sections and immunocytochemistry. Acta Neuropathol. 112, 389–404 (2006).

76. Marttinen, M. et al. A multiomic approach to characterize the temporal sequence in Alzheimer’s disease-related pathology. Neurobiol. Dis. 124, 454–468 (2019).

77. Martiskainen, H. et al. Effects of Alzheimer’s disease-associated risk loci on cerebrospinal fluid biomarkers and disease progression: a polygenic risk score approach. J. Alzheimers. Dis. 43, 565–573 (2015).

78. Natunen, T. et al. Effects of NR1H3 genetic variation on the expression of liver X receptor α and the progression of Alzheimer’s disease. PLoS One 8, e80700 (2013).

79. Natunen, T. et al. Diabetic phenotype in mouse and humans reduces the number of microglia around β-amyloid plaques. Mol. Neurodegener. 15, 66 (2020).

80. Kemppainen, S. et al. Organotypic Hippocampal Slice Cultures from Adult Tauopathy Mice and Theragnostic Evaluation of Nanomaterial Phospho-TAU Antibody-Conjugates. Cells 12, 1422 (2023).

81. Gabbouj, S. et al. Intranasal insulin activates Akt2 signaling pathway in the hippocampus of wild-type but not in APP/PS1 Alzheimer model mice. Neurobiol. Aging 75, 98–108 (2019).

82. Hao, Y. et al. Dictionary learning for integrative, multimodal and scalable single-cell analysis. Nat. Biotechnol. 42, 293–304 (2024).

83. Choudhary, S. & Satija, R. Comparison and evaluation of statistical error models for scRNA-seq. Genome Biol. 23, (2022).

84. Korsunsky, I. et al. Fast, sensitive and accurate integration of single-cell data with Harmony. Nat. Methods 16, 1289–1296 (2019).

85. Young, M. D. & Behjati, S. SoupX removes ambient RNA contamination from droplet-based single-cell RNA sequencing data. Gigascience 9, giaa151 (2020).

86. Shan, X. et al. Fully defined NGN2 neuron protocol reveals diverse signatures of neuronal maturation. Cell reports methods 4, 100858 (2024).

87. Andreatta, M. & Carmona, S. J. UCell: Robust and scalable single-cell gene signature scoring. Comput. Struct. Biotechnol. J. 19, 3796–3798 (2021).

88. Van de Sande, B. et al. A scalable SCENIC workflow for single-cell gene regulatory network analysis. Nat. Protoc. 2020 157 15, 2247–2276 (2020).

89. Wolf, F. A., Angerer, P. & Theis, F. J. SCANPY: Large-scale single-cell gene expression data analysis. Genome Biol. 19, 15 (2018)

