## supplementary Figures for "Non-microglial downregulation of *PLCG2* impairs synaptic function and elicits Alzheimer disease-related hallmarks"

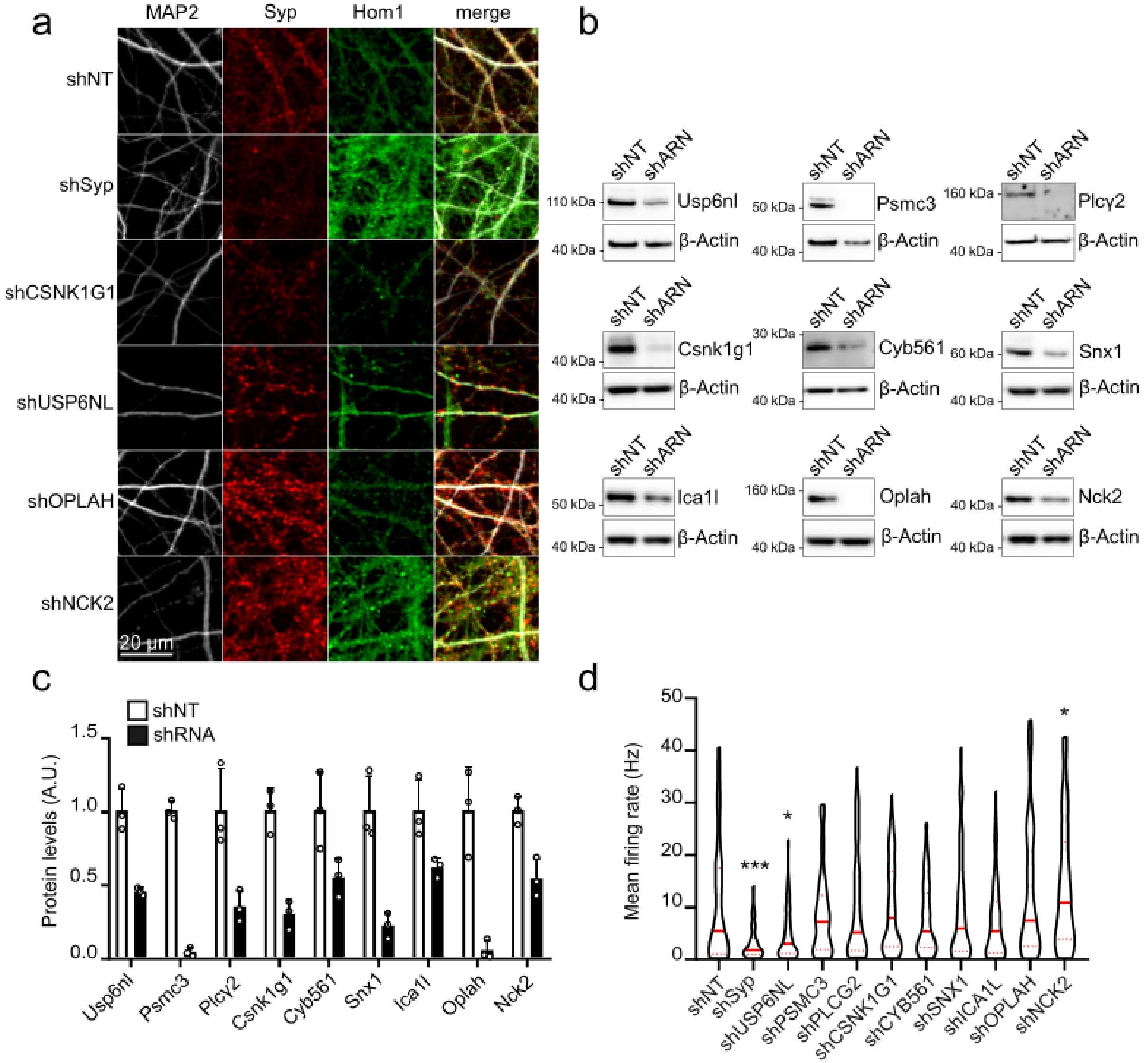


**Supplementary figure S1. (a)** Representative images of MAP2, Syp and Homer1 immunostainings following shRNA-mediated downregulation of *Syp, Csnk1g1, Usp6NL, Oplah* and *Nck2* at MOI 4. **(b)** Western blots showing the validation of gene downregulation for Usp6nl, Psmc3, Plcγ2, Csnk1g1, Cyb561, Snx1, Ica1l, Oplah, and Nck2 transduced at MOI 2 in PNCs. **(c)** Quantification of protein levels in PNCs transduced with these nine shRNAs, normalized by the non-targeting control (shNT) levels. Data given as mean±SD, n=3. **(d)** Impact of these nine shRNAs on the mean spike frequency in PNCs at DIV21, measured by MEA. Violin plots show median and quartile lines, n=3. Statistical analyses were performed using Kruskall-Wallis test followed by Dunn's multiple comparison test. *p<0.05, *** p<0.001.

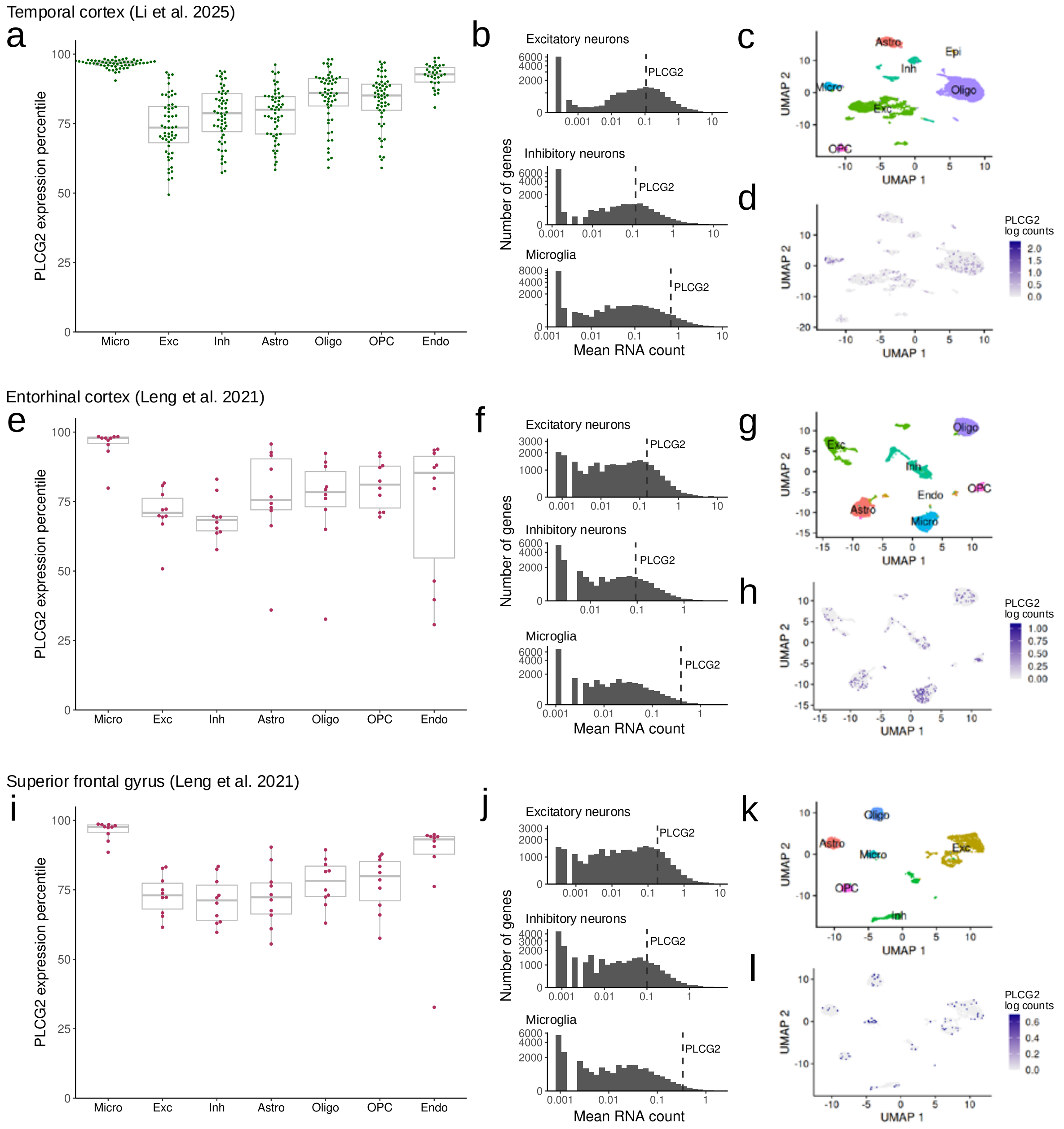


**Supplementary figure S2.** Single-cell RNA-seq datasets show that PLCG2 is expressed in human neurons. **(a)** After ranking genes by mRNA abundance in seven cell types within each subject, the percentile of PLCG2 among the rankings is shown. Data is from temporal cortex tissue[22]. **(b)** Histograms of genes' mRNA abundance in three cell types in data from a representative subject. The position of PLCG2 in the distribution is indicated. Zero values were replaced with 0.8 times the lowest non-zero value. **(c)** UMAP dimensional reduction and cell-type labeling in a representative subject. **(d)** Feature plot of PLCG2 expression in the same subject. **(e-h)** Same as above, but for entorhinal cortex (Leng et al. 2021). **(i-l)** Same as above, but for superior frontal gyrus[2386]. Micro = microglia, Exc = excitatory neurons, Inh = inhibitory neurons, Astro = astrocytes, Oligo = oligodendrocytes, OPC = oligodendrocyte precursor cells, Endo = endothelial cells.

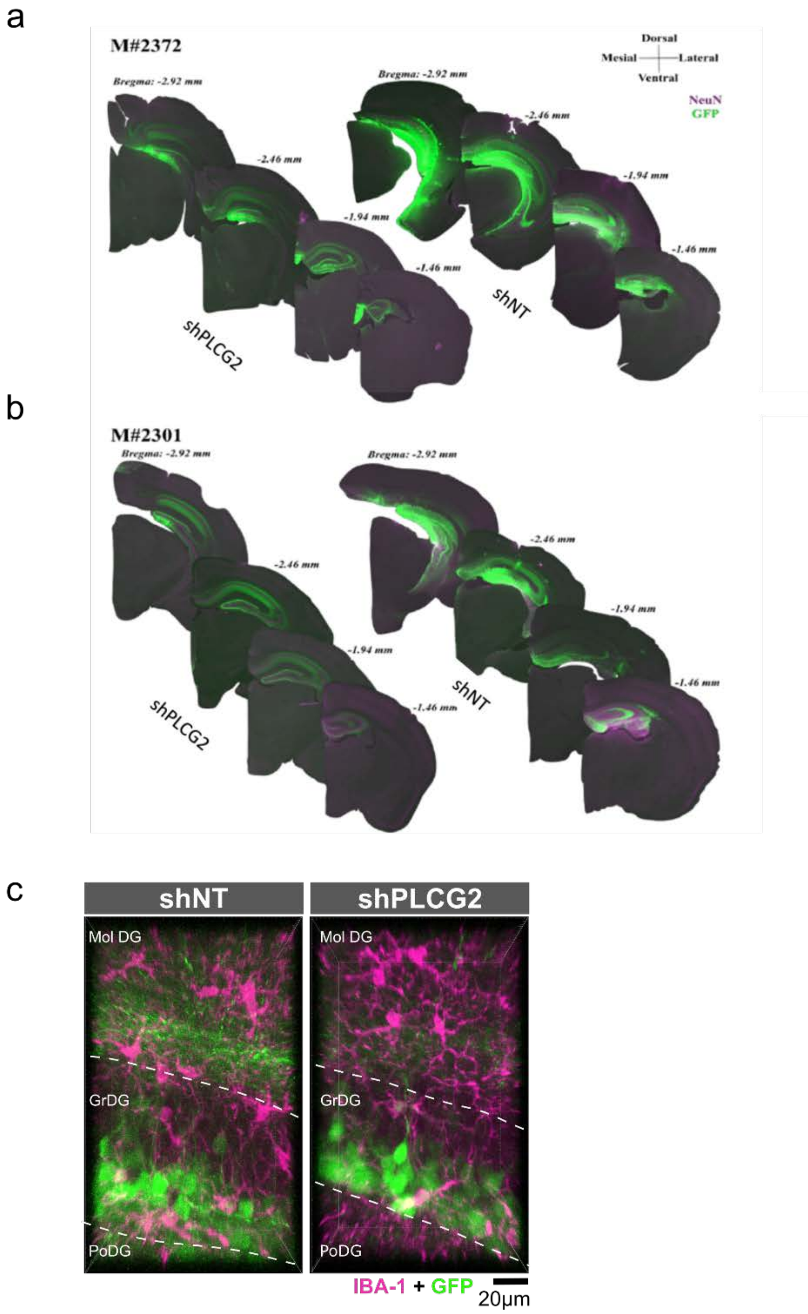


**Supplementary figure S3.** Extent of transduction by LV constructs following stereotactic injection in the hippocampus. **(a, b)** Representative images of contra-lateral injection of shNT/ shPLCG2 lentiviral vectors in the dorsal dentate gyrus in two different animals (#2372 and #2301 in panels a and b, respectively); these images were taken at 10× magnification on an epifluorescence microscope where sections were labeled with NeuN (in magenta) and GFP (in green), arranged from anterior (-1.46 µm) to posterior (-2.92 µm). **(c)** Representative images showing the absence of colocalization between transfected (GFP, in green) and microglial (IBA-1, in magenta) cells; MolDG= Molecular layer of dentate gyrus, GrDG= granular layer of dentate gyrus, PoDG= polymorph layer of dentate gyrus.

**Supplementary figure S4.** Properties of action potentials in shPLCG2 vs shNT transduced DG granule cells. **(a)** Traces of action potentials triggered by a short depolarizing pulse from -70 mV in shPLCG2 (red) vs shNT (grey) transduced DG granule cells. **(b-d)** Analysis of action potential amplitude **(b)**, threshold for action potential discharge **(c)** and action potential half-width **(d)**. Mann-Whitney tests were applied to compare the average values for shNT (n=15) *vs* shRNA (n=14) transduced DG granule cells.

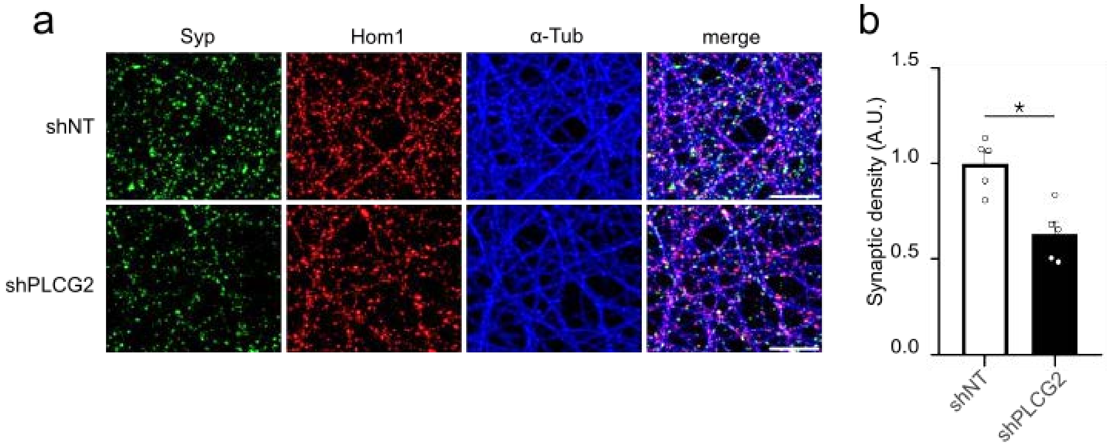


**Supplementary figure S5.** *Plcg2* downregulation decreases synaptic density in primary rat neurons cultured in microfluidic devices. **(a)** Representative immunofluorescence images of Syp, Homer1, and α tubulin markers in shNT and shPLCG2 transduced rat PNCs in microfluidic devices. **(b)** Synapse density is given as mean ± SEM, n=5 independent cultures, * p<0.05. Scale bar= 20µm.

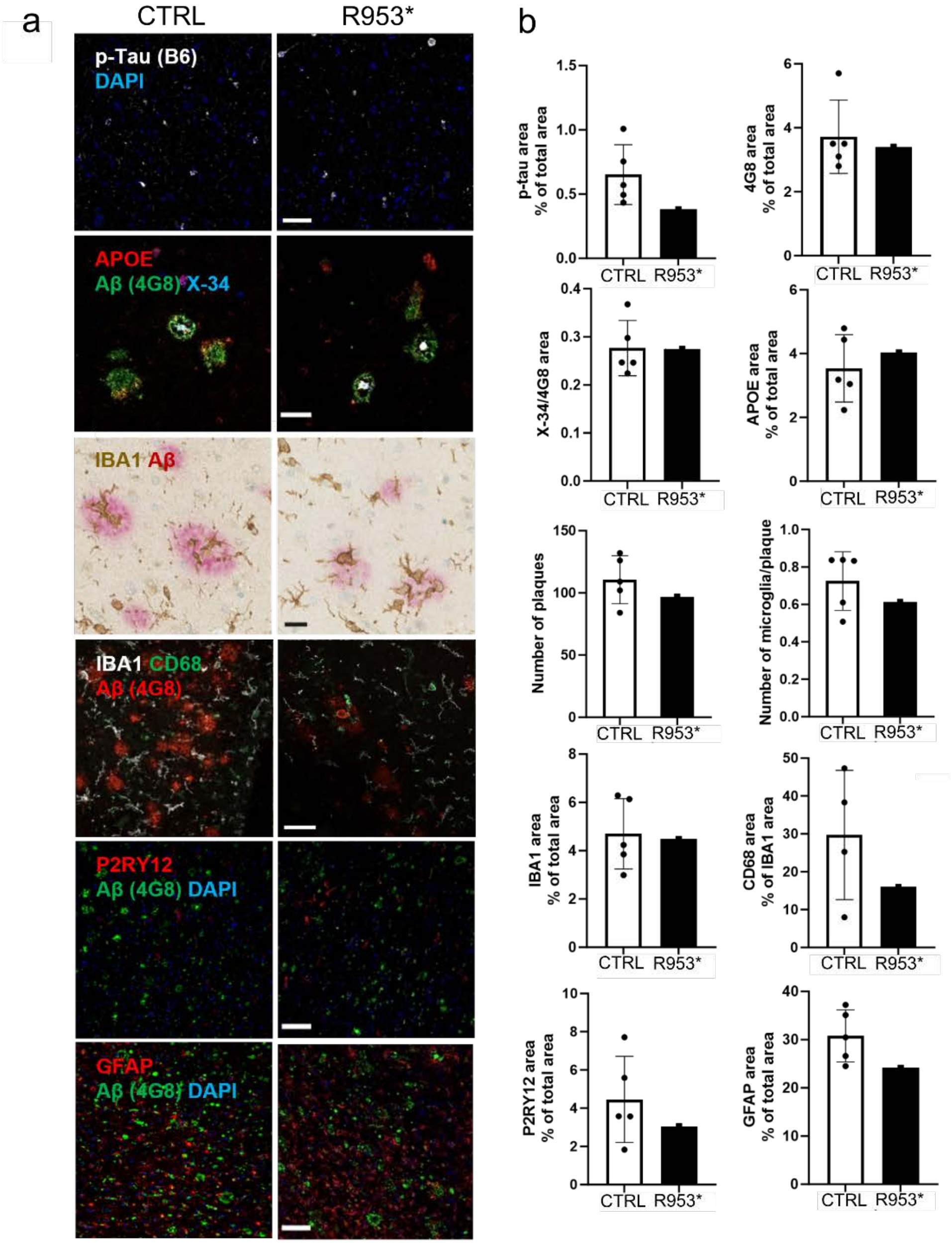


**Supplementary figure S6. (a)** Representative images of brain samples from CTRL and p.R953* carrier showing (from top to bottom): Tau phosphorylated at Ser202, Thr205 and Ser208 (p-tau, white) and nuclei (blue); APOE (red), β-amyloid plaques stained with X-34 (blue), and Aβ stained with 4G8 (green); Aβ (pink) surrounded by IBA1 positive microglia (brown); Aβ (red), IBA1 positive microglia (white) and activated microglia stained with microglial lysosomal marker CD68 (green); Aβ (green), homeostatic microglia stained with P2RY12 (red), and nuclei (blue); Aβ (green), GFAP-positive astrocytes (red), and nuclei (blue). **(b)** Bar graphs show quantification of p-tau, Aβ (4G8), APOE, IBA1, P2RY12 and GFAP immunopositive area, ratio of compact and diffuse β-amyloid plaques (X-34/4G8), total number of plaques, IBA1 positive microglia within and surrounding the plaque area, and proportion of activated microglia of IBA1 positive microglia. Scale bars = 25 µm for IBA1/Aβ, 50 µm for all the others. Data is presented as mean ± SD.

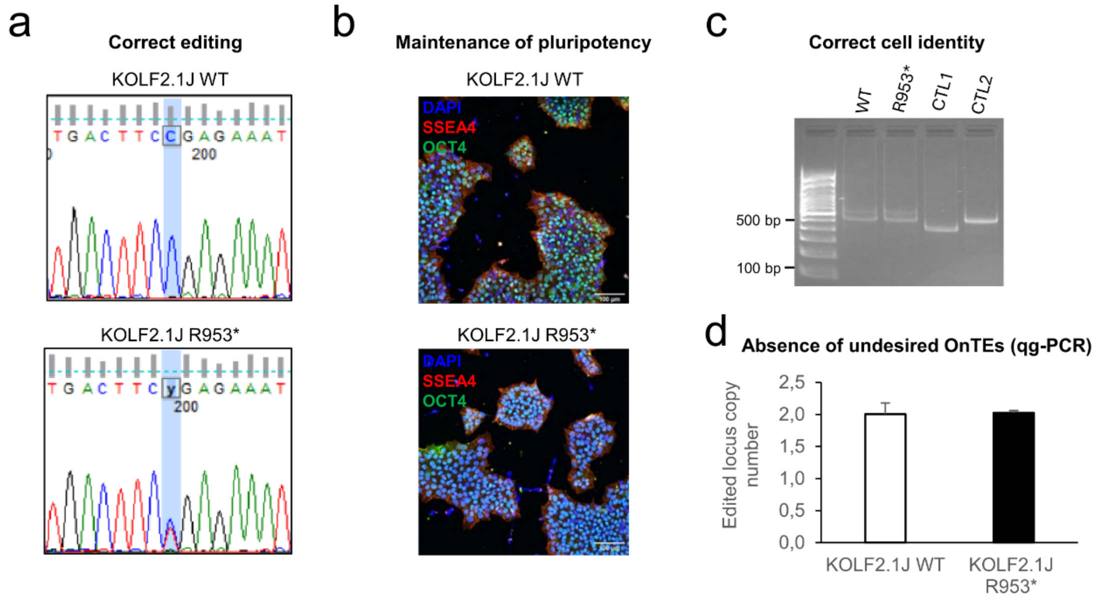


e

**Absence of undesired On TEs (WGS)**

| ID | Chrom | Pos | REF | ALT | GT | | AD | | GQ | |
|---|---|---|---|---|---|---|---|---|---|---|
| | | | | | WT | R953* | WT | R953* | WT | R953* |
| chr16:81923904:T:C | chr16 | 81923904 | T | C | 0/1 | 0/1 | 20,22 | 12,14 | 99 | 99 |
| chr16:81927346:T:A | chr16 | 81927346 | T | A | 0/1 | 0/1 | 19,21 | 19,27 | 99 | 99 |
| chr16:81935243:T:C | chr16 | 81935243 | T | C | 0/1 | 0/1 | 15,23 | 20,21 | 99 | 99 |
| chr16:81936583:C:G | chr16 | 81936583 | C | G | 0/1 | 0/1 | 11,23 | 21,23 | 99 | 99 |
| chr16:81936748:G:A | chr16 | 81936748 | G | A | 0/1 | 0/1 | 14,23 | 13,29 | 99 | 99 |
| chr16:81936761:A:C | chr16 | 81936761 | A | C | 0/1 | 0/1 | 17,23 | 13,26 | 99 | 99 |

f

**Loci screened for undesired Off TEs (WGS)**

| Chrom | Start | End | Strand | Locus | Off-target sequence | MIT Off-target Score | CFD Off-target Score |
|---|---|---|---|---|---|---|---|
| chr8 | 124094947 | 124094969 | + | exon:FER1L6 | GGGAGGATATCCCGGAAGTCGGG | 0,4371 | 0,5926 |
| chr17 | 59210228 | 59210250 | - | exon:SMG8 | GAGAGAACTTCTCGGAATTCAGG | 0,1845 | 0,5614 |
| chr17 | 50605010 | 50605032 | - | intron:CACNA1G | TAGAGAATTTCTAGGAAGTCAGG | 0,3519 | 0,3239 |
| chr10 | 22486556 | 22486578 | - | intergenic:SPAG6-PIP4K2A | GGACGGATTTCTTGGAATTCTGG | 0,1206 | 0,2800 |
| chr9 | 132845561 | 132845583 | + | intron:AK8 | GAGGGGATCACTTGGAAGTCAGG | 0,3116 | 0,2769 |
| chr17 | 4953217 | 4953239 | - | intron:ENO3 | GGTCAGATTTCTCAGAAGTCAGG | 0,2208 | 0,2600 |
| chr1 | 17888084 | 17888106 | + | intergenic:ACTL8-RP11-174G17.2 | GATGGGAGTTCACGGAAGTCAGG | 0,6941 | 0,1467 |
| chr4 | 146518038 | 146518060 | - | intron:SLC10A7 | GAGAGGATTTCCTGGAAGGCAGG | 0,0950 | 0,1396 |
| chr17 | 40825902 | 40825924 | + | intergenic:KRT10-TMEM99 | GAGAAGACTTCTGGGAAGTCAGG | 0,5537 | 0,0995 |
| chr9 | 110567804 | 110567826 | - | intron:SVEP1 | CAGGGGATGTCTAGGAAGTCTGG | 0,3554 | 0,0850 |
| chr1 | 6860681 | 6860703 | + | intron:CAMTA1 | GGGCGGATTACTTGGAGGTCAGG | 0,2173 | 0,0847 |
| chr6 | 40964279 | 40964301 | - | intergenic:RP11-570K4.1-UNC5CL | GGGGGGATTTCTTGGAAGGCTGG | 0,2001 | 0,0700 |
| chr1 | 147026072 | 147026094 | + | intergenic:RP4-704D21.2-RNVU1-8 | GATCGGATTTCTCAGAATCCAGG | 0,0146 | 0,0690 |
| chr1 | 163661134 | 163661156 | + | intergenic:RNA5SP63-U3 | GAGCACCTTTCTCTGAAGTCAGG | 0,0881 | 0,0689 |
| chr1 | 40848307 | 40848329 | - | intergenic:KCNQ4-CITED4 | GAGAGGATTTAGGGGAAGTCTGG | 0,1512 | 0,0431 |
| chr19 | 11423203 | 11423225 | - | intron:CCDC151 | GAGCGGATGGCTCCGAAGACGGG | 0,0384 | 0,0379 |
| chr14 | 87090762 | 87090784 | - | intergenic:RP11-736P16.1-RP11-1141N12.1 | GGGCTGATTTCACGGGAGTCAGG | 0,1316 | 0,0369 |
| chrY | 292953 | 292975 | - PAR1 | intron:PLCXD1 | GAGCGGAGGGCTGGGAAGTCGGG | 0,2928 | 0,0310 |
| chr2 | 59427853 | 59427875 | - | intergenic:AC007131.2-AC007179.1 | GATTGGATTTCTCTCAAGTCTGG | 0,0592 | 0,0291 |
| chr4 | 72868307 | 72868329 | - | intergenic:ADAMTS3-RNU4ATAC9P | GAGAGGAGTTGTGGGAAGTCAGG | 0,3073 | 0,0211 |
| chr12 | 132603027 | 132603049 | - | intergenic:RP13-554M15.8-LRCOL1 | GAGCGGCCTTCTGGGAAGCCTGG | 0,1252 | 0,0165 |
| chr14 | 25113974 | 25113996 | - | intergenic:STXBP6-RP11-89K22.1 | CAGCTGATTTCTCTGAAGACTGG | 0,0785 | 0,0163 |
| chr21 | 45259088 | 45259110 | + | intergenic:LINC00334-POFUT2 | GAGCGGCTTTCTCTGGAATCGGG | 0,0051 | 0,0155 |
| chr2 | 68356298 | 68356320 | + | intergenic:CNRIP1-AC015969.3 | GAGAGGATTTCTCTGGAGCCTGG | 0,0128 | 0,0119 |
| chr22 | 48341268 | 48341290 | + | intergenic:MIR3201-RP11-536P6.3 | GAGCGGATGGCTGGGAAGCCTGG | 0,0997 | 0,0116 |
| chr7 | 100091159 | 100091181 | + | intron:COPS6 | GAGGGGATTCCTTGGAAGTGGGG | 0,2396 | 0,0110 |
| chr4 | 111681920 | 111681942 | - | intergenic:RP11-255I10.2-RP11-269F21.1 | GTGGGGATTTCTCGGAACTGAGG | 0,1367 | 0,0102 |
| chr3 | 47481900 | 47481922 | + | intergenic:SCAP-ELP6 | GAGCGAATTTCTGGGAACTGTGG | 0,0298 | 0,0038 |
| chr1 | 58207616 | 58207638 | - | intergenic:RP11-393I23.4-OMA1 | GAGGGGATTTCTCGAATGTGGGG | 0,0694 | 0,0037 |
| chr11 | 61364332 | 61364354 | + | exon:TMEM138 | GAGCGGTTTTCTGGGAACTGTGG | 0,0330 | 0,0017 |
| chr13 | 114058320 | 114058342 | + | intron:RASA3 | GAGCGGATTCCTCGGGCGTGAGG | 0,0370 | 0,0011 |
| chr16 | 65062283 | 65062305 | - | intron:CDH11 | AAGCGGATTTCTTGGTTGTCTGG | 0,0413 | 0,0000 |
| chr7 | 49095781 | 49095803 | + | intergenic:AC096752.1-AC010971.1 | GAGTGCATTTCTCAGCAGTCGGG | 0,0233 | 0,000, |

g

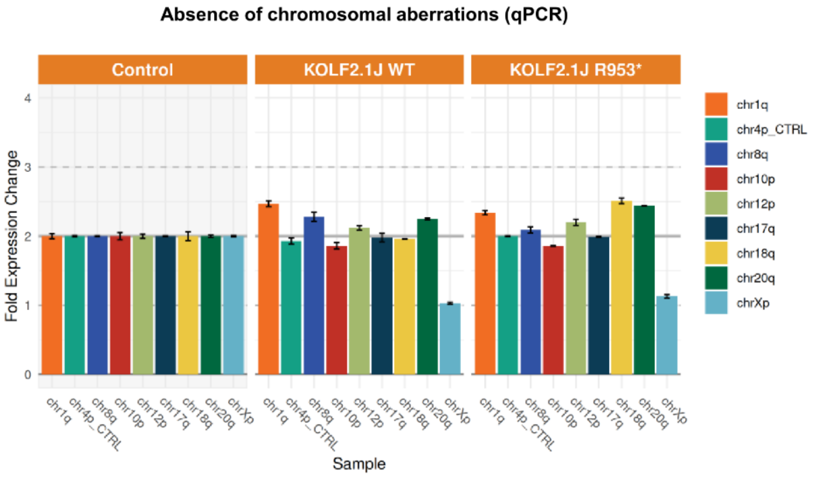


h

**Absence of aneuploidy (WGS)**

| Chromosome | Ploidy | | Mean coverage | |
|---|---|---|---|---|
| | WT | R953* | WT | R953* |
| chr1 | 2.10 | 2.08 | 36.70 | 36.94 |
| chr2 | 2.03 | 2.05 | 35.50 | 36.28 |
| chr3 | 2.05 | 2.07 | 35.84 | 36.76 |
| chr4 | 1.94 | 1.99 | 33.92 | 35.30 |
| chr5 | 2.02 | 2.05 | 35.28 | 36.26 |
| chr6 | 2.02 | 2.05 | 35.32 | 36.32 |
| chr7 | 2.04 | 2.04 | 35.64 | 36.12 |
| chr8 | 2.01 | 2.02 | 35.00 | 35.80 |
| chr9 | 2.07 | 2.06 | 36.06 | 36.46 |
| chr10 | 2.06 | 2.04 | 35.92 | 36.18 |
| chr11 | 2.09 | 2.08 | 36.44 | 36.86 |
| chr12 | 2.07 | 2.06 | 36.04 | 36.48 |
| chr13 | 1.93 | 1.98 | 33.66 | 35.16 |
| chr14 | 2.07 | 2.07 | 36.12 | 36.60 |
| chr15 | 2.15 | 2.11 | 37.48 | 37.36 |
| chr16 | 2.11 | 2.04 | 36.90 | 36.12 |
| chr17 | 2.17 | 2.07 | 37.94 | 36.76 |
| chr18 | 1.98 | 2.01 | 34.62 | 35.68 |
| chr19 | 2.16 | 1.99 | 37.64 | 35.28 |
| chr20 | 2.15 | 2.10 | 37.58 | 37.16 |
| chr21 | 1.97 | 1.99 | 34.34 | 35.28 |
| chr22 | 2.25 | 2.11 | 39.34 | 37.44 |
| chrX | 1.02 | 1.03 | 17.80 | 18.22 |
| chrY | 0.92 | 0.94 | 16.12 | 16.64 |

**Supplementary figure S7. Quality control of the KOLF2.1J iPSCs, CRISPR-edited for the PLCG2 R953* variant**. **(a)** Control for the correct editing of the KOLF2.1J iPSCs (WT) for the PLCG2 R953* variant using Sanger sequencing. **(b)** Control of iPSCs pluripotency was confirmed by immunofluorescence of OCT4 and SSEA4 markers in both WT and R953* edited iPSCs. Scale bars = 100 µm. **(c)** Fingerprinting via PCR genotyping of a microvariation at the human D1S80 locus on chromosome 1 confirmed the similar identity between PLCG2 WT and PLCG2 R953* iPSCs compared to controls 1 and 2 (CTL1, human genomic DNA; CTL2, HEK293 cells genomic DNA). **(d, e)** The absence of undesired on-target effects (e.g. loss of heterozygosity or complex rearrangement) was confirmed using qPCR **(d)** and heterozygous SNP genotyping of the edited PLCG2 locus using WGS **(e)**. ID: identifier of the variant composed of chr:position:reference_allele:alternate_allele according to the GRCh38 assembly; Chrom: chromosome according to the GRCh38 assembly; Pos: position according to the GRCh38 assembly; REF: reference allele; ALT: alternative allele; GT: genotype, 0: reference allele, 1: alternate allele; AD: allelic depths for the reference and the alternate alleles in the order listed; GQ: genotype quality, which represents the Phred-Scaled confidence that the genotype assignment is correct. The value of GQ is capped at 99. **(f)** List of potential off-target loci that were predicted by CRISPOR and checked by WGS. WGS analysis confirmed the absence of off-target effects of the KOLF2.1 WT iPSC genome edition to generate the R953* variant. Quantitative PCR **(g)** and WGS **(h)** analyses confirmed the absence of CRISPR-induced chromosomal aberration or aneuploidy.

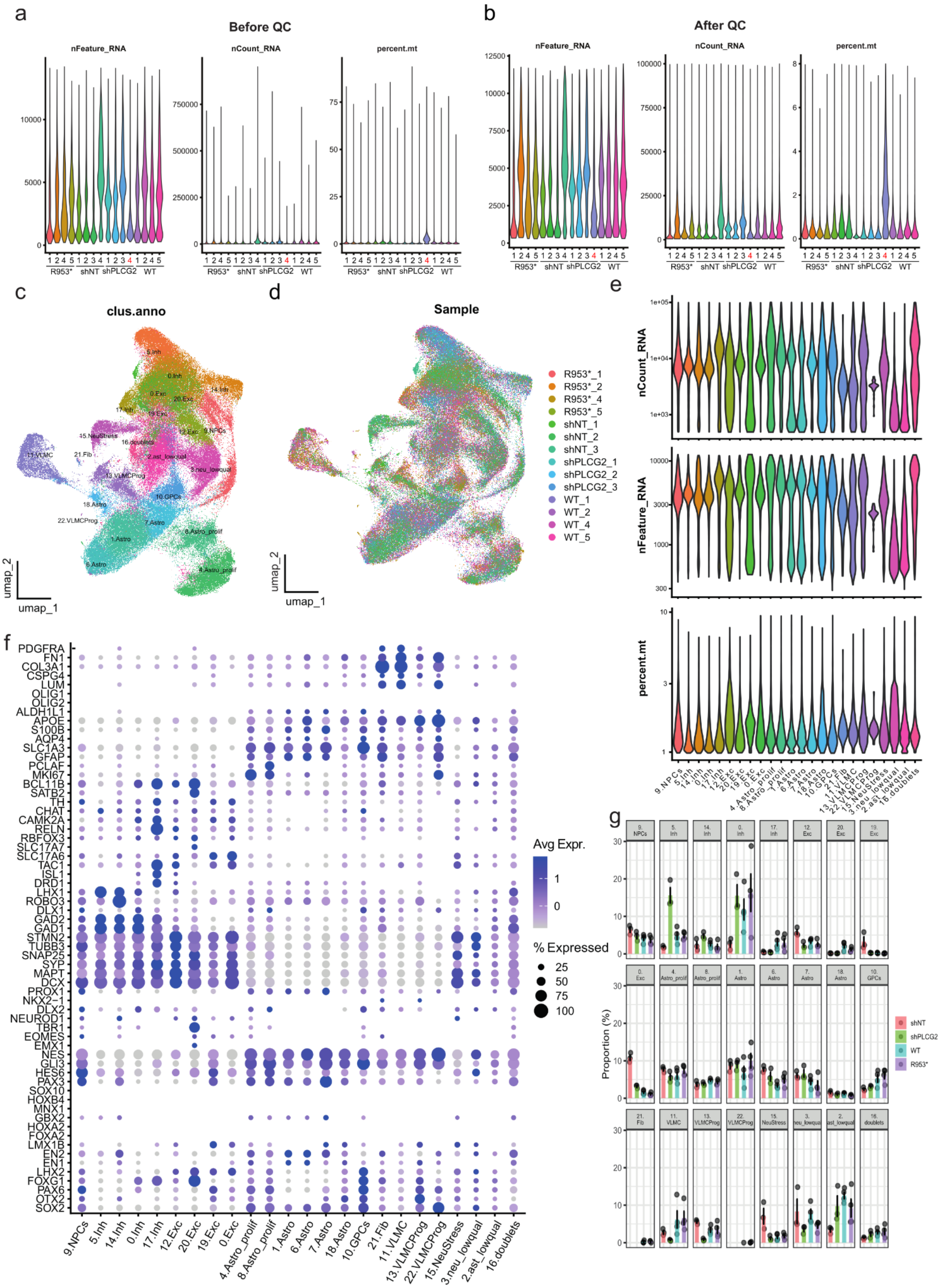
a
Before QC
nFeature_RNA
nCount_RNA
percent.mt
R953*
shNT
shPLCG2
WT
b
After QC
c
clus.anno
umap_1
umap_2
d
Sample
R953*_1
R953*_2
R953*_4
R953*_5
shNT_1
shNT_2
shNT_3
shPLCG2_1
shPLCG2_2
shPLCG2_3
WT_1
WT_2
WT_4
WT_5
e
f
PDGFRA
FN1
COL3A1
CSPG4
LUM
OLIG1
OLIG2
ALDH1L1
APOE
S100B
AQP4
SLC1A3
GFAP
PCLAF
MKI67
BCL11B
SATB2
TH
CHAT
CAMK2A
RELN
RBFOX3
SLC17A7
SLC17A6
TAC1
ISL1
DRD1
LHX1
ROBO3
DLX1
GAD2
GAD1
STMN2
TUBB3
SNAP25
SYP
MAPT
DCX
PROX1
NKX2−1
DLX2
NEUROD1
TBR1
EOMES
EMX1
NES
GLI3
HES6
PAX3
SOX10
HOXB4
MNX1
GBX2
HOXA2
FOXA2
LMX1B
EN2
EN1
LHX2
FOXG1
PAX6
OTX2
SOX2
9.NPCs
5.Inh
14.Inh
0.Inh
17.Inh
12.Exc
20.Exc
19.Exc
0.Exc
4.Astro_prolif
8.Astro_prolif
1.Astro
6.Astro
7.Astro
18.Astro
10.GPCs
21.Fib
11.VLMC
13.VLMCProg
22.VLMCProg
15.NeuStress
3.neu_lowqual
2.ast_lowqual
16.doublets
Avg Expr.
% Expressed
25
50
75
100
g
Proportion (%)
shNT
shPLCG2
WT
R953*

**Supplementary figure S8. (a-b)** Violin plots of QC metrics of the nuclei analyzed in each sample (nFeature_RNA: number of genes detected, nCount_RNA: number of RNA, percent.mt: percentage of mitochondrial RNA), before QC **(a)** and after the first stage of QC **(b)**. Low quality sample highlighted in red have been excluded from downstream analyses. **(c-d)** UMAP of first stage QC'ed nuclei clustered according to their transcriptome similarity colored by cluster annotation **(c)** or sample origin **(d). (e)** Violin plots of QC metrics for each cluster. **(f)** Dot plot of expression of the neuronal cell type markers in each cluster. **(g)** Proportions of each cluster in the samples.

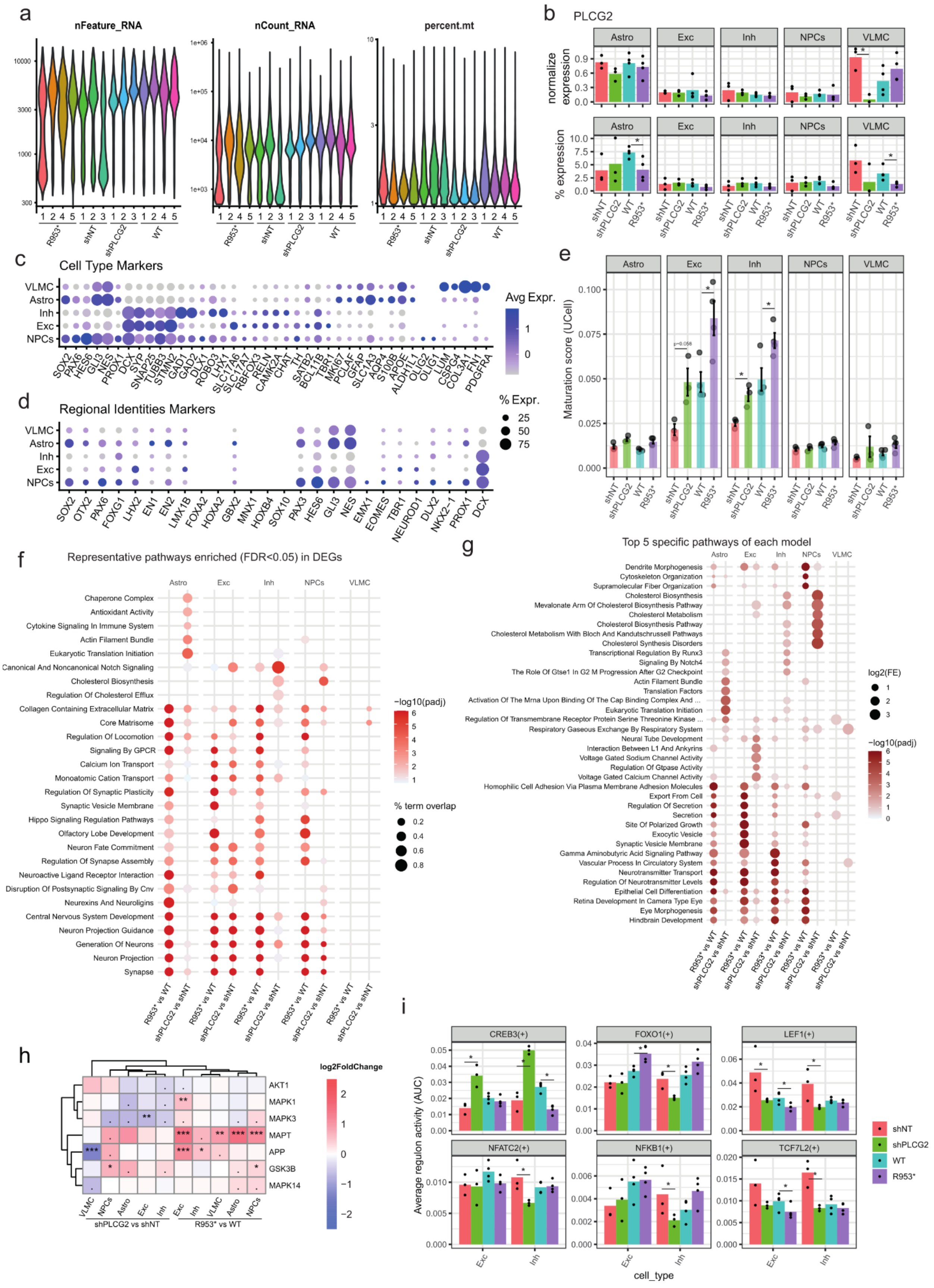

a
nFeature_RNA
nCount_RNA
percent.mt
R953*
shNT
shPLCG2
WT
b
PLCG2
Astro
Exc
Inh
NPCs
VLMC
normalize expression
% expression
shNT
shPLCG2
WT
R953*
c
Cell Type Markers
Avg Expr.
% Expr.
d
Regional Identities Markers
e
Maturation score (UCell)
f
Representative pathways enriched (FDR<0.05) in DEGs
Chaperone Complex
Antioxidant Activity
Cytokine Signaling In Immune System
Actin Filament Bundle
Eukaryotic Translation Initiation
Canonical And Noncanonical Notch Signaling
Cholesterol Biosynthesis
Regulation Of Cholesterol Efflux
Collagen Containing Extracellular Matrix
Core Matrisome
Regulation Of Locomotion
Signaling By GPCR
Calcium Ion Transport
Monoatomic Cation Transport
Regulation Of Synaptic Plasticity
Synaptic Vesicle Membrane
Hippo Signaling Regulation Pathways
Olfactory Lobe Development
Neuron Fate Commitment
Regulation Of Synapse Assembly
Neuroactive Ligand Receptor Interaction
Disruption Of Postsynaptic Signaling By Cnv
Neurexins And Neuroligins
Central Nervous System Development
Neuron Projection Guidance
Generation Of Neurons
Neuron Projection
Synapse
−log10(padj)
% term overlap
R953* vs WT
shPLCG2 vs shNT
g
Top 5 specific pathways of each model
log2(FE)
−log10(padj)
h
log2FoldChange
AKT1
MAPK1
MAPK3
MAPT
APP
GSK3B
MAPK14
shPLCG2 vs shNT
R953* vs WT
i
CREB3(+)
FOXO1(+)
LEF1(+)
NFATC2(+)
NFKB1(+)
TCF7L2(+)
Average regulon activity (AUC)
cell_type

**Supplementary Figure S9: (a)** Violin plots of number of genes, number of RNA and mitochondrial RNA percentage detected in nuclei for each sample after final QC. **(b)** Mean PLCG2 expression or percentage detection in each cell type. **(c-d)** Dot plot of the expression in different cell types **(c)** and regional identities markers **(d)**. **(e)** Average maturation score by cell type and sample, using UCell with the 25 genes of the neuronal maturation associated signature (MS-NS) from Shan et al. (2024)[86] **(f)** Pathways enriched in the DEGs of each PLCG2 downregulation model for each cell type. Representative pathways selected from the top100 MSigDB CP and GO gene-sets enriched (FDR<0.05, hypergeometric test) in each cell type and models. **(g)** Top five pathways per cell type exclusively enriched in one of the two PLCG2 downregulation models. **(h)** Heat map of differential expression for selected genes, tested using DESeq2. **(i)** Regulons activity score for transcription factors known to be regulated by GSK3β. TF Regulons identification using pyscenic, and regulons activity score using AUCell, statistical comparison between condition using Wilcoxon rank-sum test. *: p-value<0.1.

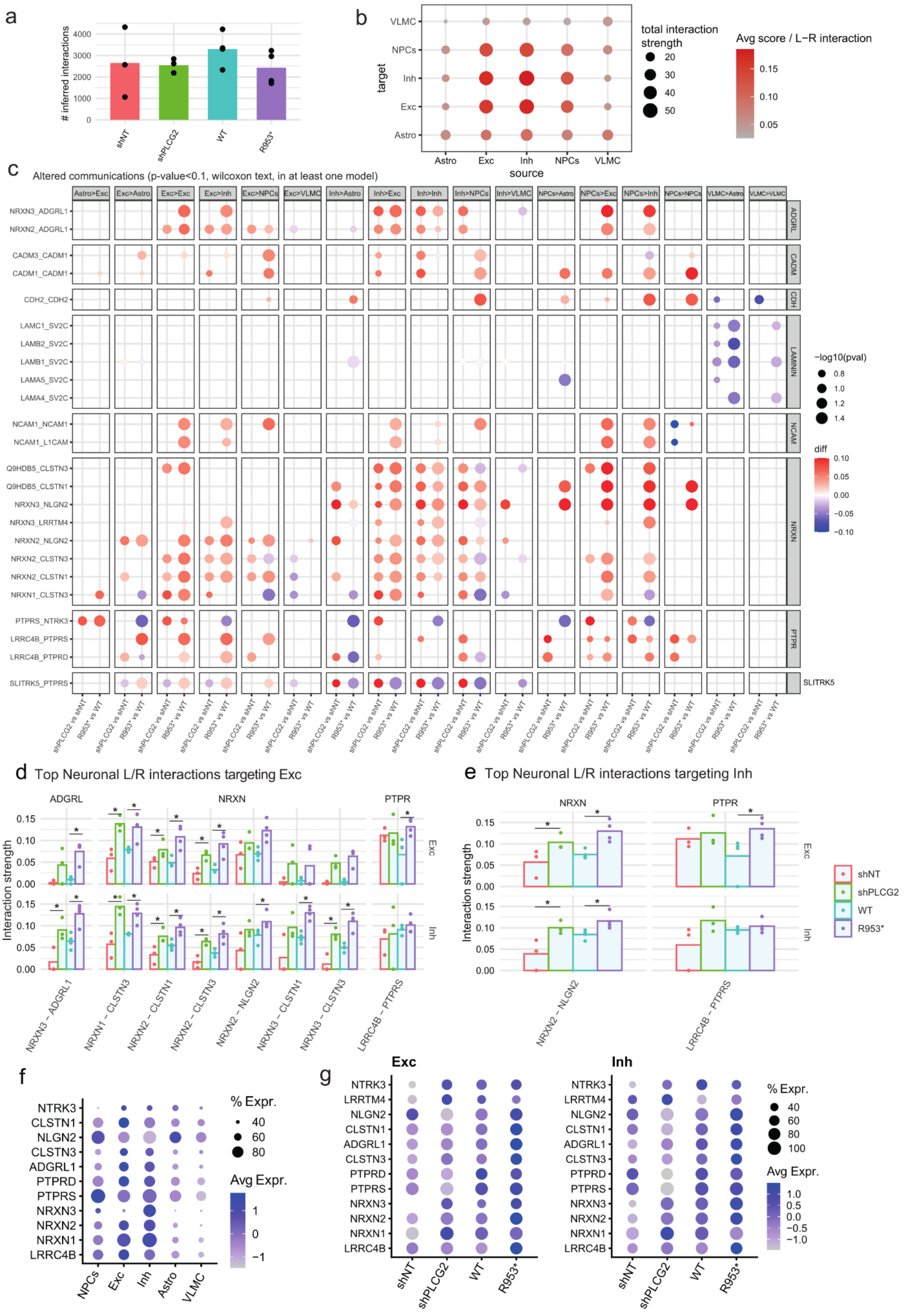
a
### inferred interactions
shNT
shPLCG2
WT
R953*
b
target
source
total interaction strength
Avg score / L−R interaction
c
Altered communications (p-value<0.1, wilcoxon text, in at least one model)
−log10(pval)
diff
d
Top Neuronal L/R interactions targeting Exc
e
Top Neuronal L/R interactions targeting Inh
Interaction strength
f
% Expr.
Avg Expr.
g
Exc
Inh

**Supplementary Figure S10:** Cell Cell Communication using CellChat. **(a)** Number of inferred interactions per sample. **(b)** Average number of inferred paracrine interactions and overall strength between cell types. **(c)** All ligand/receptor (L/R) interactions found affected (p-value<0.1, Wilcoxon rank sum test) in at least one PLCG2 downregulation model. **(d-e)** Inferred L/R interaction strength of ADGRL, NRXN and PTPR pathways in excitatory neurons **(d)** or in inhibitory neurons **(e)**. *: p-value<0.1, Wilcoxon rank sum test. **(f-g)** Average expression for genes of top L/R signaling affected in PLCG2 reduction models by cell type **(f)** or by condition in excitatory or inhibitory neurons **(g)**.
